# The LCLAT1/LYCAT acyltransferase is required for EGF-mediated phosphatidylinositol-3,4,5-trisphosphate generation and Akt signalling

**DOI:** 10.1101/2023.01.26.524308

**Authors:** Victoria Chan, Cristina Camardi, Kai Zhang, Laura A. Orofiamma, Karen E. Anderson, Jafarul Hoque, Leslie N. Bone, Yasmin Awadeh, Daniel K. C. Lee, Norman J. Fu, Jonathan T. S. Chow, Leonardo Salmena, Len R. Stephens, Phillip T. Hawkins, Costin N. Antonescu, Roberto J. Botelho

## Abstract

Receptor tyrosine kinases such as epidermal growth factor receptor (EGFR) stimulate phosphoinositide 3-kinases (PI3Ks) to convert phosphatidylinositol-4,5-bisphosophate [PtdIns(4,5)P_2_] into phosphatidylinositol-3,4,5-trisphosphate [PtdIns(3,4,5)P_3_]. PtdIns(3,4,5)P_3_ then remodels actin and gene expression, and boosts cell survival and proliferation. PtdIns(3,4,5)P_3_ partly achieves these functions by triggering activation of the kinase Akt, which phosphorylates targets like Tsc2 and GSK3β. Consequently, unchecked upregulation of PtdIns(3,4,5)P_3_-Akt signalling promotes tumour progression. Interestingly, 50-70% of PtdIns and PtdInsPs have stearate and arachidonate at *sn*-1 and *sn*-2 positions of glycerol, respectively, forming a species known as 38:4-PtdIns/PtdInsPs. LCLAT1 and MBOAT7 acyltransferases partly enrich PtdIns in this acyl format. We previously showed that disruption of LCLAT1 lowered PtdIns(4,5)P_2_ levels and perturbed endocytosis and endocytic trafficking. However, the role of LCLAT1 in receptor tyrosine kinase and PtdIns(3,4,5)P_3_ signaling was not explored. Here, we show that LCLAT1 silencing in MDA-MB-231 and ARPE-19 cells abated the levels of PtdIns(3,4,5)P_3_ in response to EGF signalling. Importantly, LCLAT1-silenced cells were also impaired for EGF-driven and insulin-driven Akt activation and downstream signalling. Thus, our work provides first evidence that the LCLAT1 acyltransferase is required for receptor tyrosine kinase signalling.

## Introduction

Phosphoinositide (PtdInsP) lipid signaling orchestrates a variety of cellular functions such as organelle identity and membrane trafficking, ion channel activity, cytoskeletal organization, regulation of gene expression, modulation of metabolic activity, and cell proliferation and survival (Balla, 2013; Choy et al., 2017; Dickson and Hille, 2019; Doumane et al., 2022; Idevall-Hagren and De Camilli, 2015; Posor et al., 2022). PtdInsPs are generated by the reversible phosphorylation of the phosphatidylinositol (PtdIns) headgroup by several types of lipid kinases and phosphatases. Collectively, and based on the headgroup phosphorylation, these enzymes can generate up to seven species of PtdInsPs (Balla, 2013; Choy et al., 2017; Dickson and Hille, 2019; Posor et al., 2022). Nonetheless, there is another facet of PtdInsP biology that is poorly defined at the regulatory and functional levels – the control and function of the acyl composition of PtdInsPs (Barneda et al., 2019; Bozelli and Epand, 2019; Choy et al., 2017; D’Souza and Epand, 2014; Traynor-Kaplan et al., 2017).

In many mammalian tissues and cells, 50-70% of PtdIns and PtdInsPs are enriched for stearate and arachidonate at the *sn*-1 and *sn*-2 positions, respectively – this acyl combination is referred to as 38:4-PtdIns or 38:4-PtdInsPs (Anderson et al., 2013; Anderson et al., 2016; Barneda et al., 2019; D’Souza and Epand, 2014; Haag et al., 2012; Imae et al., 2012; Lee et al., 2012; Milne et al., 2005; Traynor-Kaplan et al., 2017). Additionally, this acyl composition is unique to PtdIns and PtdInsPs since other phospholipids have distinct acyl profiles (Barneda et al., 2019; Bozelli and Epand, 2019; Hicks et al., 2006; Traynor-Kaplan et al., 2017). This suggests that the acyl groups of PtdIns and PtdInsPs do more than simply embedding the lipids into the membrane bilayer (Barneda et al., 2019; Bozelli and Epand, 2019; Choy et al., 2017).

However, the exact acyl profile of PtdIns and PtdInsPs can vary between cell types, environmental conditions, and patho-physiological conditions such as cancer (Anderson et al., 2016, 10; Barneda et al., 2019; Bozelli and Epand, 2019; Hicks et al., 2006; Imae et al., 2012; Mujalli et al., 2018; Traynor-Kaplan et al., 2017). For example, p53^-/-^ cancer cells, prostate cancer cells and triple negative breast cancer cells all have distinct acyl profiles of PtdIns relative to normal tissue (Freyr Eiriksson et al., 2020; Koizumi et al., 2019; Naguib et al., 2015; Rueda-Rincon et al., 2015). Yet, there is much to be understood about the regulatory mechanisms that establish and remodel the acyl profile of PtdIns and PtdInsPs, and their functional implications.

The LCLAT1 acyltransferase has been identified as one of the enzymes that remodels and enriches PtdIns and/or PtdInsPs in stearate at the *sn*-1 position (Bone et al., 2017; D’Souza and Epand, 2014; Imae et al., 2012; Zhang et al., 2023). LCLAT1 is an ER-localized protein and is thought to act during the Lands’ Cycle or PtdIns Cycle to enrich PtdIns in stearate at *sn*-1 (Barneda et al., 2019; Blunsom and Cockcroft, 2020; Bone et al., 2017; Imae et al., 2012). Murine tissues deleted for LCLAT1 had reduced levels of 38:4-PtdIns, mono-PtdInsP, and bis-phosphorylated PtdInsPs (Imae et al., 2012). More recently, we observed that LCLAT1 silencing reduced the relative levels of endosomal phosphatidylinositol-3-phosphate [PtdIns(3)P] and phosphatidylinositol-4,5-bisphosphate [PtdIns(4,5)P_2_] on the plasma membrane, while the levels of phosphatidylinositol-4-phosphate [PtdIns(4)P] remained unchanged (Bone et al., 2017). Importantly, PtdIns and bis-phosphorylated PtdInsPs (mostly PtdIns(4,5)P_2_), but not mono-phosphorylated PtdInsPs (mostly PtdIns(4)P] were altered in their acyl profile upon LCLAT1 silencing. This effect on specific pools of PtdInsPs has also been observed in cells disrupted for LPIAT1/MBOAT7, an acyltransferase thought to enrich PtdIns/PtdInsPs in arachidonic acid (Anderson et al., 2013).

PtdIns(4,5)P_2_ regulates a number of functions including endocytosis, ion transport, and the cytoskeleton organization (Balla, 2013; Katan and Cockcroft, 2020; Sun et al., 2013). PtdIns(4,5)P_2_ is also a precursor for other signalling intermediates regulated by growth factor receptors like the Epidermal Growth Factor (EGF) and its receptor, the EGF receptor (EGFR), a major receptor tyrosine kinase (Katan and Cockcroft, 2020; Orofiamma et al., 2022). EGF binds and dimerizes EGFR, leading to receptor autophosphorylation on various tyrosine residues on the receptor’s C-terminal tail region (Böni-Schnetzler and Pilch, 1987; Gullick et al., 1985; Honegger et al., 1987; Koland and Cerione, 1988; Linggi and Carpenter, 2006; Yarden and Schlessinger, 1987). Motifs harbouring these phospho-tyrosines serve as docking sites for adaptor proteins like Grb2, which assemble a signaling complex composed of other protein kinases and phosphatases, and lipid-metabolizing enzymes (Holgado-Madruga et al., 1996, 2; Margolis et al., 1990a; Orofiamma et al., 2022; Rodrigues et al., 2000). For example, active EGFR recruits and activates Phospholipase Cγ (PLCγ), which hydrolyses PtdIns(4,5)P_2_ into diacylglycerol and inositol-1,4,5-trisphosphate (IP_3_), and which releases Ca^2+^ from endoplasmic reticulum stores (Delos Santos et al., 2017; Margolis et al., 1990a; Margolis et al., 1990b). In addition, Gab1, recruited to the membrane via interactions with EGFR-bound Grb2, engages Class I PI3Ks to convert PtdIns(4,5)P_2_ to PtdIns(3,4,5)P_3_ (Kiyatkin et al., 2006; Rodrigues et al., 2000). This burst of PtdIns(3,4,5)P_3_ then recruits and activates PDK1 and Akt protein kinases (Alessi et al., 1997; Bellacosa et al., 1998; Manning and Toker, 2017; Stokoe et al., 1997). Akt is a major driver of cell metabolism and growth by phosphorylating numerous targets like GSK3β and TSC2 (Cross et al., 1995; Inoki et al., 2002; Manning and Toker, 2017; Rodgers et al., 2017; Sugiyama et al., 2019). For example, Akt inactivates TSC2, a GTPase activating protein (GAP) for the Rheb GTPase, thus promoting the mTORC1 pathway (Inoki et al., 2002; Inoki et al., 2003). The PtdIns(3,4,5)P_3_-Akt-mTORC1 pathway is a major driver of cell growth, proliferation, survival, and differentation (Dey et al., 2017; Dibble and Manning, 2013; Manning and Toker, 2017; Sugiyama et al., 2019). As a result, mutations that hyperactivate this pathway are often associated with human cancers like triple-negative breast cancer (TNBC) (Dey et al., 2017; Li et al., 2017).

Overall, given that LCLAT1 is a PI acyltransferase and the connection between EGF and PI3K-Akt pathway, we postulated that LCLAT1 disruption would impair EGF-mediated PtdIns(3,4,5)P_3_-Akt signalling, reflecting a role for PI acyl profile specificity for signaling by this pathway. In fact, we found that LCLAT1 silencing in at least two cell lines impeded generation of PtdIns(3,4,5)P_3_ and Akt activation upon addition of EGF.

## Methods and Materials

### Cell Culture

The male human-derived ARPE-19 retinal pigment epithelial cell line was obtained from ATCC (CRL-2302, Manassas, VA) and was cultured in DMEM/F12 medium (ThermoFisher Scientific, Mississauga, ON) supplemented with 10% fetal bovine serum (Wisent, St. Bruno, QB), 100 U/mL penicillin and 100 µg/mL streptomycin (ThermoFisher Scientific). The female human-derived MDA-MB-231 triple negative breast cancer cell line was obtained from ATCC (CRM-HTB-26). Wild-type MDA-MB-231 cells and its derivatives (see below) were cultured in DMEM medium (Wisent) supplemented with 10% fetal bovine serum, 100 U/mL penicillin and 100 µg/mL streptomycin. The female human-derived HEK293T cell line was cultured in DMEM with 10% fetal calf serum and 1% penicillin/streptomycin. All cells were cultured at 37°C and 5% CO_2_. *Mycoplasma* screening is performed at least annually.

### Transfection and siRNA-mediated gene silencing

To silence gene expression of LCLAT1 in both ARPE-19 and MDA-MB-231 cells, custom-synthesized siRNA oligonucleotides against LCLAT1 were designed using Horizon Discovery siDESIGN Centre. We designed and tested siLCLAT1-1, siLCLAT1-2, and siLCLAT1-3 with the respective sequences 5’-GGAAAUGGAAGGAUGACAAUU-3’, 5’CAGCAAGUCUCGAAGUAAUU-3’, and 5’-UCGAAGACAUGAUUGAUUAUU-3’.

Synthesis was by Sigma-Aldrich (Oakville, ON). In addition, we used siLCLAT1-5, a siGenome-validated oligonucleotide from Horizon (cat. D-010307-01-0002). Moreover, a non-targeting control siRNA (NT siRNA, or siCON) with the sequence 5’-CGUACUGCUUGCGAUACGGUU-3’ was used (Sigma-Aldrich). Cells were transfected with 22 pmol of siRNA oligonucleotides/well using Lipofectamine RNAiMAX (ThermoFisher Scientifc) in Opti-MEM reduced serum media (ThermoFisher Scientific) for 3 h at 37°C and 5% CO_2_ as per manufacturer’s instructions. After transfection, cells were washed and incubated with fresh growth medium. Two rounds of transfection were performed, 72 and 48 h before each experiment.

### Plasmids and transfections

Plasmid encoding Akt-PH-GFP was a kind gift from the Balla lab, NIH (Addgene: #51465) and was previously described in (Várnai and Balla, 1998). Akt-PH-GFP plasmid was transfected into ARPE-19 and MDA-MB-231 cells using Lipofectamine 3000 as instructed by the manufacturer. The plasmid encoding eGFP-PLCδ-PH was previously described in (Stauffer et al., 1998) and used to generate MDA-MB-231 cells stably expressing the PtdIns(4,5)P_2_ biosensor.

### Generation of doxycycline-inducible expression of eGFP-PLCδ-PH in MDA-MB-231 cell

A plasmid based on the Sleeping Beauty pSBtet-BP vector (GenScript, Piscataway, NJ; catalog number: SC1692) for inducible expression of eGFP-PLCδ1-PH was generated by gene synthesis of the open reading frame of eGFP-PLCδ-PH and insertion into the NheI and ClaI restriction sites of the pSBtet-BP vector, as described previously (Zak and Antonescu, 2023). MDA-MB-231 cells were transfected with the engineered pSBtet-BP::eGFP-PLCδ-PH and pCMV(CAT)T7-SB100 plasmid (Addgene, Plasmid #34879) using FuGENE HD transfection reagent (Promega) as instructed by manufacturer. After transfection, cells were washed and incubated with fresh growth medium for another 24 h to let cells recover in a 6-well plate, and then transferred into a T75 flask in the presence of 3 μg/mL of puromycin. Growth medium with puromycin was replaced every 2-3 days for 3 weeks. After selection, cells were treated with doxycycline (100-200 nM) for 24 h to detect eGFP-PLCδ-PH expression by Western blotting and fluorescence microscopy.

### EGF signaling and Western blotting

Before lysate preparation, cells were incubated with 2 mL of serum free growth medium for 1 h. After serum starvation, cells were stimulated with 5 ng/mL EGF for 5 or 10 min or left unstimulated (basal). Alternatively, cells were stimulated with 10 ng/mL insulin for 5 min. Following serum starvation and subsequent EGF stimulation, whole cell lysates were prepared in 200 µL 2x Laemmli Sample Buffer (0.5M Tris, pH 6.8, glycerol and 10% sodium dodecyl sulfate (SDS)) supplemented with a protease and phosphatase inhibitor cocktail (Complete 1x protease inhibitor (Sigma-Aldrich), 1 mM sodium orthovanadate, and 10 nM okadaic acid).

Lysates were heated at 65°C for 15 min and passed through a 27-gauge needle 10 times. Finally, 10% ß-mercaptoethanol and 5% bromophenol blue were added to cell lysates.

Proteins were resolved by Tris-glycine SDS-PAGE and transferred on to a polyvinylidene difluoride (PVDF) membrane. The PVDF membrane was blocked for 1 h at room temperature in blocking buffer composed of 3% bovine serum albumin (BSA) in 1X final Tris-buffered saline-Tween (TBS-T; 20 nM Tris, 150 mM NaCl and 0.1% Tween-20). After blocking, membranes were washed 3 times with wash buffer and incubated with 1:1000 primary antibody overnight at 4°C. The next day, membranes were washed 3 times, for 5 min each, and then subjected to 1:1000 secondary antibody for 1 h at room temperature. After incubation with secondary antibody, membranes were washed 3 times, 5 min each, and imaged using the ChemiDoc Imaging System (BioRad). Membranes were exposed to Immobilon Crescendo Western HRP substrate (Millipore Sigma) for 30-60 s and chemiluminescent images were acquired by the ChemiDoc System. Western blot signals were analyzed and quantified using the ImageLab 6.1 (BioRad). Band intensity was obtained by signal integration in an area corresponding to the appropriate band. This value was then normalized to the loading control signal. For quantifying phosphorylated protein levels, the phosphorylated protein signal and the corresponding total protein signal were first normalized to their respective loading controls, followed by the ratio of corrected phosphorylated protein signal to total protein signal.

Primary antibodies raised in rabbit were anti-LCLAT1 (cat. 106759, GeneTex), anti-phospho-Akt (S473, cat. 9271), anti-phospho-Akt1 (S473, cat. 9018), anti-phospho-Akt2 (S474, cat. 8599), anti-Akt1 (cat. 2938), anti-phospho-EGFR (Y1068. Cat. 2234), anti-phospho-tuberin/TSC2 (T1462, cat. 3611), anti-tuberin/TSC2 (cat. 3612), anti-phospho-GSK3ß (S9, cat. 9323), anti-phospho-ERK1/2 (T202/Y204, monoclonal, cat. 9201), anti-ERK1/2 (monoclonal, 137F5, cat. 4695), anti-p53 (monoclonal, 7F5, cat. 2527), anti-p21 Waf1/Cip1 (monoclonal, 12D1, cat. 2947), anti-cofilin (monoclonal, D3F9), anti-vinculin (polyclonal, Cat. 4550) anti-clathrin, (monoclonal DC36, cat. 4796), and anti-GAPDH (cat. 2118) were all from Cell Signaling Technologies. Antibodies raised in mouse were anti-Akt (monoclonal, 40D4; cat. 2920), anti-Akt2 (cat. 5239), anti-GSK3ß (cat. 9832), and anti-puromycin (cat. MABE343) and were all from Cell Signaling Technology. Goat anti-EGFR antibodies were from Santa Cruz Biotechnology (sc-03-G). Horseradish peroxidase (HRP)-linked secondary anti-rabbit, anti-mouse, and anti-goat IgG antibodies were from Cell Signaling Technology.

### Fluorescence and Immunofluorescence

To detect surface levels of EGFR in MDA-MB-231 and ARPE-19 cells, cells were blocked with 3% BSA in PBS supplemented with 1 mM CaCl_2_ and MgCl_2_ for 30 min on ice, followed by 1 h incubation with a 1:200 dilution of mouse anti-EGFR antibody collected in-house from the mAb108 hybridoma obtained from ATCC (Cabral-Dias et al., 2022). After washing with PBS, cells were fixed with 4% paraformaldehyde in PBS for 15 min and then quenched with 100 mM glycine in PBS for 10 min, followed by washing with PBS, and then fluorescent secondary mouse antibodies (Jackson ImmunoResearch Labs Inc., West Grove, PA) at a 1:500 dilution in 1% BSA in PBS for 1 h at room temperature. The coverslips were mounted using Dako fluorescence mounting medium (Agilent Technologies, Inc. Mississauga, ON, Canada). For labeling the plasma membrane, cells with stained with 7 µg/mL FM4-64FX and imaged within 10 min to minimize internalization of FM4-64FX.

### Microscopy

Confocal and TIRF micrographs were obtained using a Quorum Diskovery spinning disc confocal system coupled to a TIRF module (Quorum Technologies, Inc., Guelph, ON). The microscope itself consisted of an inverted fluorescence microscope (DMi8; Leica) equipped with an Andor Zyla 4.2 Megapixel sCMOS camera (Oxford Instruments, Belfast, UK), a 63x oil immersion objective (1.4 NA), and standard excitation and emission filter sets and lasers were used for all fluorophores. The TIRF module used a 63x/NA1.49 objective with a 1.8x camera relay (total magnification 108x). The microscope system was controlled by the MetaMorph acquisition software (Molecular Devices, LLC, San Jose, CA, USA).

For fixed cells, z-stacks of 10-30 images were acquired with an inter-planal distance of 0.6 µm distance. For live-cell imaging of Akt-PH-GFP dynamics in ARPE-19 cells or of eGFP-PLCδ-PH in MDA-MB-231 cells by TIRF microscopy or spinning disc confocal respectively, cells were maintained in DMEM free of phenol red or serum in a Chamlide microscope-mounted chamber at 37°C and 5% CO_2_. For timelapse of Akt-PH-GFP in MDA-MB-231 cells, a baseline was obtained by acquiring images for 1 min at 15 s, then EGF was added, and images acquired every 15 s for 10 min.

### Image Analysis and Processing

Image processing and quantitative analysis were performed using ImageJ or FIJI v. 2.3 (Schindelin et al., 2012) or Volocity v. 7 (Quorum Technologies), where image enhancements were completed without altering the quantitative relationship between image elements. For quantification of EGFR cell surface, regions of interest were generated by freehand to define the cell outline, the mean fluorescence intensity over the whole cell area was calculated, and then background corrected (Cabral-Dias et al., 2022). Mean fluorescence from at least 30 cells per condition per experiment was then normalized against control condition. To obtain the relative cell surface localization index for the Akt-PH-GFP probe in ARPE-19 cells, we used ImageJ to determine the ratio of TIRF/epifluorescence fluorescence for >100 cells per condition per experiment. To measure the plasma membrane to cytosolic ratio of eGFP-PLCδ-PH in MDA-MB-231 cells (Cabral-Dias et al., 2021), we first defined the plasma membrane using the FM4-64FX channel to randomly selected regions of the plasma membrane and cytosol for each cell and their ratio was calculated. For Akt-PH-GFP timelapse movies were acquired and randomly selected regions in the cell periphery and cytosol were selected in FIJI and then the plasma membrane to cytosol fluorescence ratio over time. We examined at least 23 cells over three experiments.

### Lipid extraction

Cells were grown to ∼0.5×10^6^ per well in a 6-well plate. Cells were then placed on ice and the media removed. Cells were scraped in 500 μL of ice cold 1M HCl and transferred to a pre-cooled safe lock 2 mL microcentrifuge tube. Cells were then collected by centrifugation at 13,000 xg for 10 min at 4°C. The supernatant was aspirated, and the pellet was then snap frozen in liquid nitrogen.

### Mass Spectrometry Lipid Analysis

Mass spectrometry was used to measure PtdInsPs from lipid extracts prepared from 0.5-1×10^6^ MDA-MB-231 cells or 0.6×10^6^ ARPE-19 cells as previously described (Clark et al., 2011). Briefly, we used a QTRAP 4000 mass spectrometer (AB Sciex, Macclesfield, UK) and employing the lipid extraction and derivatization method described for cultured cells (Clark et al., 2011). The response ratios of a specific PtdInsP acyl species were calculated by normalizing the targeted lipid integrated response area to that of a known amount of added relevant internal standard. A ratio of ratios was then calculated by taking the response ratios of 38:4-PtdIns, 38-4-mono-PtdInsP, 38-4-bis-PtdInsP, and 38:4-PtdIns(3,4,5)P_3_ against the sum of response ratios for 36:1 and 36:2 (36:x) of PtdIns or the corresponding 36:x-mono-PtdInsP or 36:x-bis-PtdInsP. Data are presented as mean ± STD from four separate experiments.

### Statistical analysis

Experiments were repeated a minimum of three independent times, with the exact number for each experiment indicated in the respective figure legend and/or graph as individual data points. Microscopy data were selected and quantified randomly, i.e. before inspection of cells. If regions of a cell were selected, this was done with the independent channel prior to quantification of the target channel. Data were collected as mean ± standard deviation (STD) or ± standard error of the mean (SEM). Statistical comparisons of means were then performed with GraphPad Prism v. 10. Statistical tests were selected based on data conditions such as number of parameters, sample size, assumption of normality, number of comparisons made, and correction for multiple comparisons. Figure legends specify the tests employed for a given data set such as one-sample t-test, one-way or repeated-measures two-way ANOVA tests, and recommended post-hoc tests. p values are shown, where p< 0.05 was typically accepted as significantly different.

## Results

### LCLAT1 acyltransferase regulation of EGFR trafficking

In our previous work, LCLAT1 silencing altered the endocytosis and endosomal trafficking of the transferrin receptor in ARPE-19 cells (Bone et al., 2017). We thus queried if LCLAT1 disruption would alter the total and surface levels of EGFR and/or affect EGFR signalling in at least two human-derived cells lines: the non-cancerous, male-derived ARPE-19 and the female-derived triple negative breast cancer cell line, MDA-MB-231. To do this, we used previously designed and validated siRNA oligonucleotides against LCLAT1 (Bone et al., 2017) and a non-targeting oligonucleotide (see Methods), transiently transfected cells twice, and after 48 h, we lysed the cells and probed for LCLAT1 levels by Western blotting. As shown in Figure 1A and Figure 2A respectively, control ARPE-19 and MDA-MB-231 cells displayed a major band at 35 kDa. After transfection with siLCLAT1-1, this band was reduced by ∼70% intensity in both cell types, manifesting efficient LCLAT1 silencing (Fig. 1A, B and Fig. 2A, B). We also observed reduced LCLAT1 expression in ARPE-19 and MDA-MB-231 cells transfected with independent siRNA oligonucleotides against LCLAT1 (Supplemental Figure S1A, 1B, S2A, 2B, and S4).

**Figure 1:**
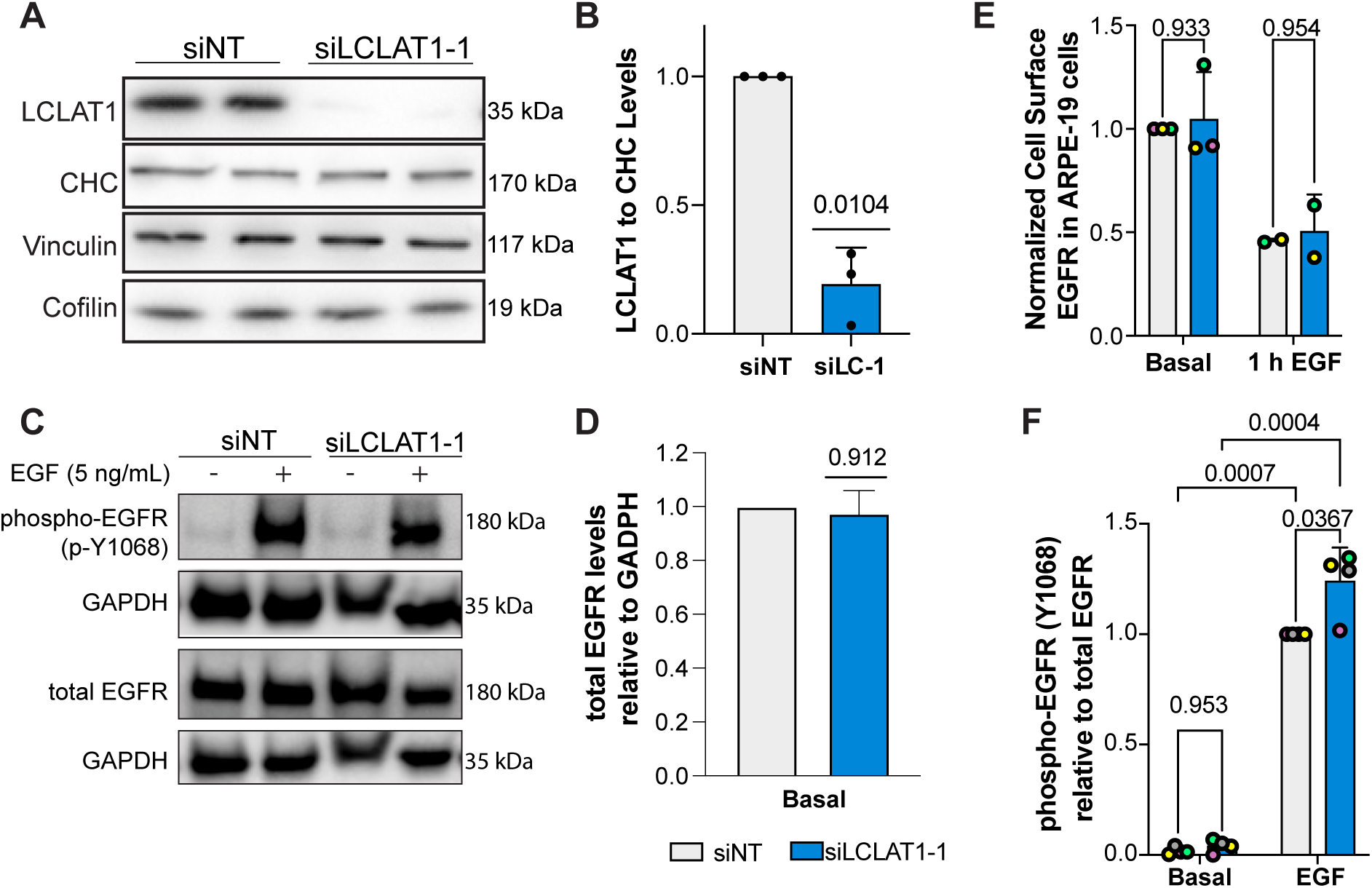
LCLAT1 silencing has no negative impact on EGFR activation, EGFR total levels, and EGFR surface levels in ARPE-19 cells. **A.** Western blots showing repressed LCLAT1 expression in ARPE-19 cells transfected with siLCLAT1-1 oligonucleotides relative to non-targeting control siRNA (NT siRNA). Two replicate lanes per condition are shown. Clathrin heavy chain (CHC), cofilin, and vinculin were used as loading controls. **B.** Normalized ratio of LCLAT1 expression to CHC in ARPE-19 cells. **C.** ARPE-19 cells silenced for LCLAT1 or treated with non-targeting oligonucleotide were serum-starved and then stimulated with 5 ng/mL EGF for 5 min. Lysates were then probed for total EGFR or phospho-EGFR. GAPDH was employed as the loading control. **D.** Quantification of total EGFR relative to the respective GAPDH signal. **E.** Normalized cell surface EGFR detected by immunofluorescence before and after 1 h stimulation with 100 ng/mL EGF in non-targeted and LCLAT1-silenced ARPE-19 cells. **F.** Quantification of phospho-EGFR (p-Y1068) relative to the respective total EGFR signal. All experiments were repeated a minimum of three times except in E (EGF stimulation, n =2). Data points from matching independent experiments are colour coded. For B, D, and F shown is mean ±STD. Data in E is shown as mean ± SEM where at least 40-80 cells were scored per condition per experiment. Data in B and D were analysed by a one-sample t-test using hypothetical value of 1. For data in E and F, a repeated measures two-way ANOVA followed by Sidak’s (E) or Tukey’s (F) post-hoc test was used. *p* values are indicated.

**Figure 2:**
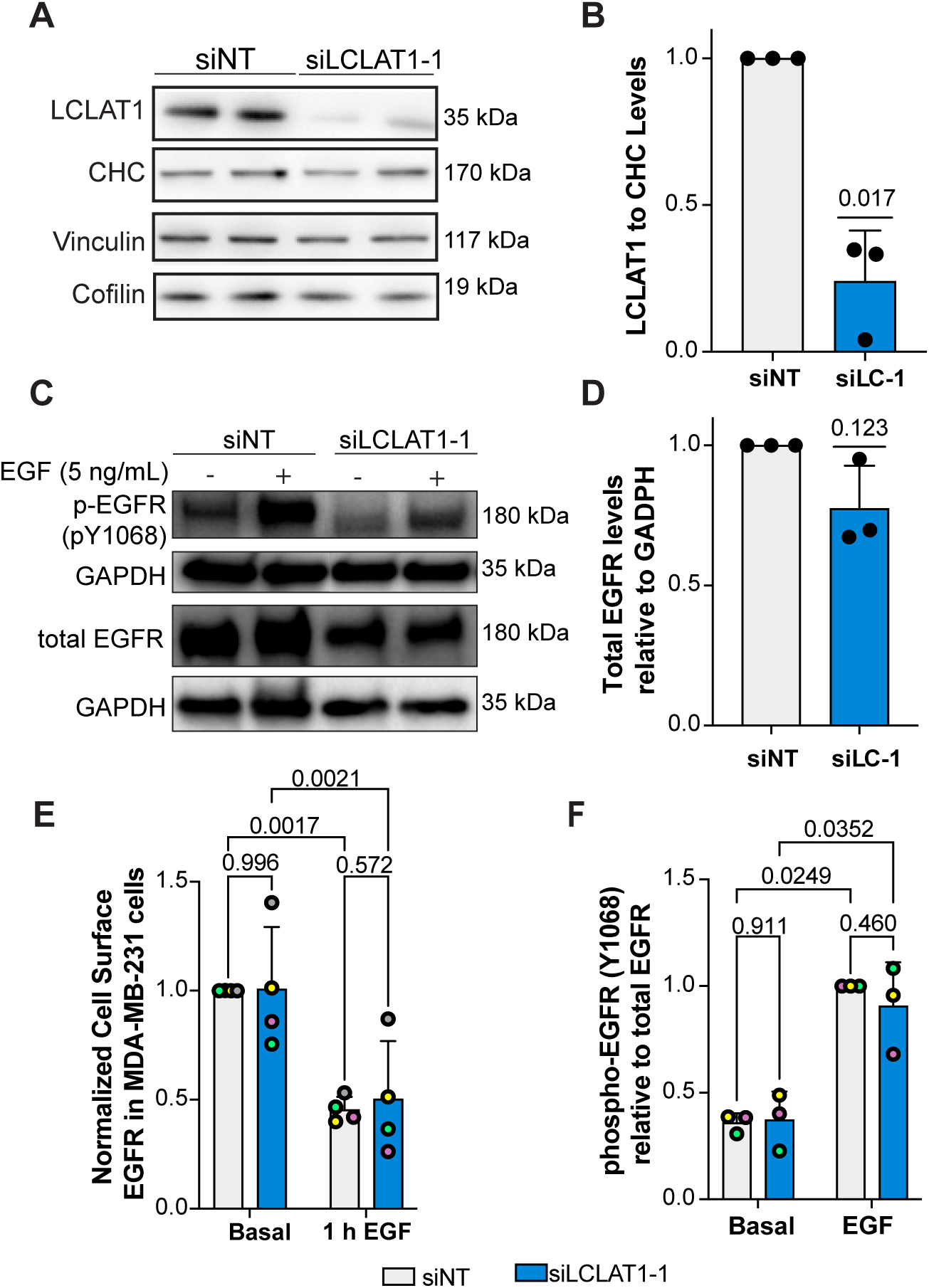
LCLAT1 silencing has no negative impact on EGFR activation, total EGFR levels, and surface EGFR levels in MDA-MB-231 cells. **A.** Western blot showing LCLAT1 silencing in MDA-MB-231 cells transfected with siLCLAT1-1 (siLC-1) relative to non-targeting control siRNA (NT siRNA). Each conditions shows two replicates. Clathrin heavy chain (CHC), vinculin, and cofilin were used as a loading control. **B.** Normalized ratio of LCLAT1 expression to CHC signal in MDA-MB-231 cells. **C.** MDA-MB-231 cells silenced for LCLAT1 or treated with non-targeting oligonucleotides were serum-starved and then stimulated with 5 ng/mL EGF for 5 min. Lysates were then probed for total EGFR or phospho-EGFR (pY-1068). GAPDH was employed as the loading control. **D.** Quantification of total EGFR relative to respective GAPDH. **E.** Normalized cell surface EGFR detected by immunofluorescence before and after 1 h stimulation with 100 ng/mL EGF in non-targeted and LCLAT1-silenced MDA-MB-231 cells. **F.** Quantification of phospho-Y1068-EGFR relative to EGFR. All experiments were repeated at least three independent times. Data points from matching independent experiments are colour coded. For B, D, and F, shown are the mean ± STD. For E, the mean ± SEM is shown where 50-100 cells were scored per condition per experiment. Data in B and D were analysed by a one-sample t-test using hypothetical value of 1. For data in E and F, a repeated measures two-way ANOVA followed by Tukey’s post-hoc test was used. p values are shown.

We next examined the effects of LCLAT1 silencing on total and surface levels of EGFR in these cells. We observed that LCLAT1 silencing in ARPE-19 and MDA-MB-231 cells did not alter total EGFR levels as measured by Western blotting (Fig. 1C, 1D and 2C, 2D) nor the EGFR surface levels as measured by immunofluorescence of unpermeabilized cells under basal conditions (Fig. 1E and 2E). Moreover, the surface levels of EGFR after 1 h of 100 ng/mL EGF stimulation dropped by a similar extend in both control and LCLAT1-silenced ARPE-19 (Fig. 1E) and MDA-MB-231 (Fig. 2E). Importantly, LCLAT1-silencing did not impair EGF-mediated phosphorylation of Y1068 of EGFR in ARPE-19 cells (Fig. 1C, F) and MDA-MB-231 cells (Fig. 2C, F); in fact, EGFR phosphorylation at Y1068 appears elevated in LCLAT1-silenced cells (Fig. 2F). Overall, these data suggest that the steady-state levels and trafficking of EGFR, and immediate response to EGF in ARPE-19 and MDA-MB-231 cells did not decline during LCLAT1 silencing.

### LCLAT1 acyltransferase silencing reduces PtdIns(3,4,5)P_3_ synthesis in response to EGF

We previously observed that ARPE-19 cells silenced for LCLAT1 had ∼30% reduction in PtdIns(4,5)P_2_ levels (Bone et al., 2017). To determine if this was recapitulated in MDA-MB-231 cells, we generated cells stably engineered for doxycycline-inducible eGFP-PLCδ-PH, a reporter for PtdIns(4,5)P_2_ (Stauffer et al., 1998). We then quantified the fluorescence ratio of eGFP-PLCδ-PH on the plasma membrane over cytosolic signal by using FM4-64FX to define the plasma membrane. Upon silencing of LCLAT1 in these cells, the eGFP-PLCδ-PH fluorescence ratio of plasma membrane to cytosol declined significantly relative to non-silenced cells (Fig. 3A, B) suggesting that LCLAT1-silenced MDA-MB-231 cells also had less PtdIns(4,5)P_2_.

**Figure 3.**
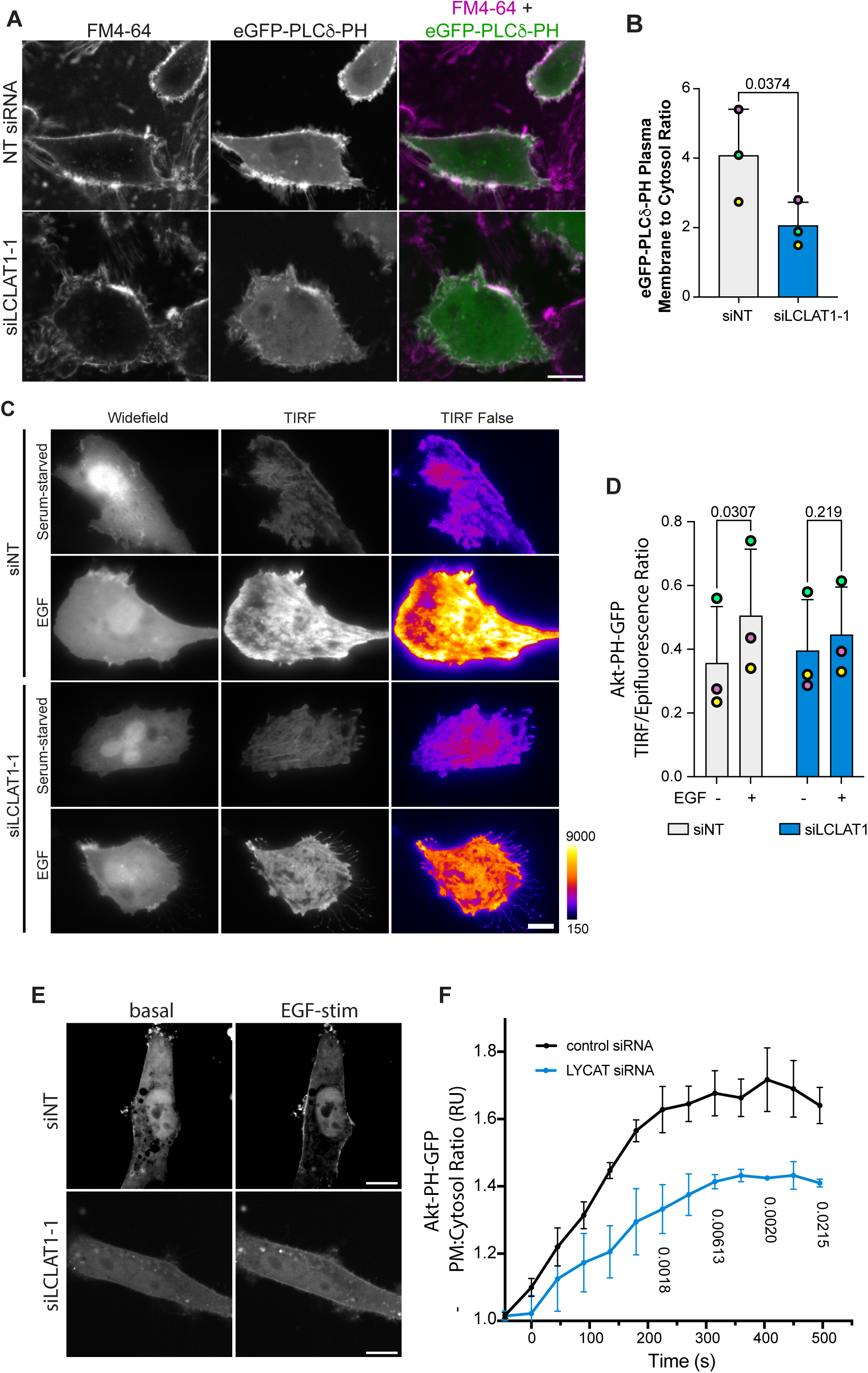
Defective PI(4,5)P_2_ and EGF-stimulated PtdIns(3,4,5)P_3_ synthesis in LCLAT1-silenced cells. MDA-MB-231 cells (A, E), and ARPE-19 (C) were mock-silenced or LCLAT1-silenced. **A.** Confocal images of MDA-MB-231 cells stably expressing eGFP-PLCδ-PH (green) and labelled with FM4-46FX (magenta). **B.** Quantification of eGFP-PLCδ-PH fluorescence on FM4-64X labelled cell periphery relative to its cytosolic signal. **C.** TIRF and epifluorescence microscopy of ARPE-19 cells mock-silenced or silenced for LCLAT1 and expressing Akt-PH-GFP. Cells were maintained serum starved or exposed to 20 ng/mL EGF for 5 min. The TIRF field is shown both in grayscale and as false-colour (fire LUT), where black-indigo is weakest and yellow-white is strongest. For representation, cells selected expressed similar levels of Akt-PH-GFP. **D.** Quantification of TIRF/epifluorescence ratio of Akt-PH-GFP. Total Akt-PH-GFP fluorescence in TIRF field is expressed as a ratio against the corresponding total fluorescence in the epifluorescence field. Shown is the ratio for serum-starved and EGF-stimulated control and LCLAT1-silenced cells. **E.** Spinning disc confocal images of MDA-MB-231 cells mock-silenced or LCLAT1-silenced before and during 20 ng/mL EGF stimulation. **F.** Quantification of Akt-PH-GFP fluorescence at the plasma membrane relative to its cytosolic signal from time-lapse imaging over 10 min of stimulation with 20 ng/mL EGF. Scale bar = 20 µm. All experiments were repeated three independent times. For B and D, data points from matching independent experiments are colour coded. Shown is the mean ± SEM, where data in D were binned every 45 sec (three images). For B and D, data are based on 30-50 transfected cells per condition per experiment. For F, a total of 23 transfected cells were traced over time over three independent experiments. Data in B was analysed by paired Student’s t-test. For D and F, repeated measures two-way ANOVA and Sidak’s post-hoc test was used to test data in D. p-values are shown.

Next, a key outcome of EGFR stimulation is the activation of PI3Ks to convert PtdIns(4,5)P_2_ to PtdIns(3,4,5)P_3_ (Hu et al., 1992; Orofiamma et al., 2022; Rodrigues et al., 2000). To determine if PtdIns(3,4,5)P_3_ synthesis was affected in LCLAT1-disturbed cells, we transfected ARPE-19 and MDA-MB-231 cells with plasmids encoding Akt-PH-GFP, a biosensor for 3-phosphorylated PtdInsPs (Várnai and Balla, 1998). The recruitment of Akt-PH-GFP to the plasma membrane in response to EGF was then quantified using two different methods. Given that ARPE-19 cells are exceptionally flat, we quantified the ratio of TIRF to epifluorescence fields (TIRF/Epi fluorescence ratio) as an indicator for PtdIns(3,4,5)P_3_ levels at the plasma membrane. While control cells readily increased their Akt-PH-GFP on the plasma membrane after EGF stimulation, cells perturbed for LCLAT1 expression displayed substantially lower TIRF/Epifluorescence of their Akt-PH-GFP (Fig. 3C, D). We then assessed if MDA-MB-231 cells were also impaired for PI3K signalling. Since these cells are rounder, we measured Akt-PH-GFP on the plasma membrane more readily in optical sections obtained from the middle of the cell by spinning disc confocal microscopy by sampling Akt-PH-GFP at the cell periphery against cytosolic signal. We tracked the Akt-PH-GFP recruitment over 10 min after adding EGF. We observed an increase in Akt-PH-GFP to the cell periphery after EGF stimulation of non-silenced MDA-MB-231 cells (Fig. 3E, F). Importantly, this increase in Akt-PH-GFP at the cell periphery was suppressed in the LCLAT1-silenced cell group (Fig. 3E, F). Hence, despite near normal levels of EGFR and p-EGFR, we reveal that LCLAT1 expression is required for EGF-mediated increase in PI(3,4,5)P_3_ levels in at least two cell types.

### The impact of LCLAT1 acyltransferase expression on the acyl profile of PtdInsPs

We next examined the PtdInsP acyl profile and their relative levels by mass spectrometry in control (cells maintained in medium supplemented with serum), serum-starved (medium with no serum for 1 h and not further stimulated), and EGF-stimulated for 5 min. These conditions were examined in both ARPE-19 and MDA-MB-231 cells that were subjected to non-targeting siRNA or LCLAT1-silencing siRNA oligonucleotides. For this analysis, we normalized lipid spectral counts against synthetic standards added to the samples to generate a response ratio (see methods). In addition, to correct for variation in cell input between experiments, we further normalized against an internal benchmark by comparing changes in 38:4 PtdIns, *mono*-PtdInsPs, *bis*-PtdInsPs, and PtdIns(3,4,5)P_3_ relative to standardized 36:x-PtdIns and the corresponding 36:x-mono-PtdInsP and 36:xbis-PtdInsP since these species have previously been shown to be less affected by LCLAT1 expression relative to 38:4 species (Bone et al., 2017; Imae et al., 2012). We note that our previous normalization benchmark used the 38:x PtdInsP (not just 38:4) to its respective 36:x PtdInsP and an internal benchmark was not used (Bone et al., 2017).

For ARPE-19 cells, the major acyl species of PtdIns, *mono*-, *bis*-, and *tris*-PtdInsP was 38:4, as expected. In fact, we only detected 38:4 acyl species for PtdIns(3,4,5)P_3_. We then compared each group of 38:4 PtdInsPs to the 36:x-PtdIns and the corresponding 36:x-PtdInsP. Relative to 36:x-PtdIns, we saw that the levels of 38:4-PtdIns were reduced in LCLAT1-silenced cells (Fig. 4A). In comparison, there was no significant difference in 38:4-*mono*-PtdInsPs relative to 36:x-PtdIns or 36:x-mono-PtdInsP in any treatment (Fig. 4B, Sup. Fig. S3A). However, 38:4-*bis*-PtdInsPs declined in LCLAT1-silenced cells relative to 36:x-PtdIns (Fig. 4C) and 36:x-bis-PtdInsPs (Sup. Fig. S3B). Lastly, EGF increased the levels of 38:4-PtdIns(3,4,5)P_3_ relative to 36:x-PtdIns in both serum-starved non-silenced and LCLAT1-silenced cells (Fig. 4D). However, LCLAT1-silenced cells had a significant reduction in 38:4-PtdIns(3,4,5)P_3_ levels relative to 36:x-PtdIns compared to non-targeted ARPE-19 cells (Fig. 4D). We could not quantitatively compare 38:4-PtdIns(3,4,5)P_3_ to 36:x-PtdIns(3,4,5)P_3_ since we did not detect the latter. Overall, this suggests that 38:4-PtdIns(3,4,5)P_3_ is the predominant acyl species of this phosphoinositide upon EGF stimulation and that LCLAT1 expression is required for its efficient synthesis. Overall, we reveal that ARPE-19 cells shift their acyl profile of PtdIns, *bis*-PtdInsPs, and PtdIns(3,4,5)P_3_ upon LCLAT1-disruption, but not for *mono*-PtdInsPs.

**Figure 4.**
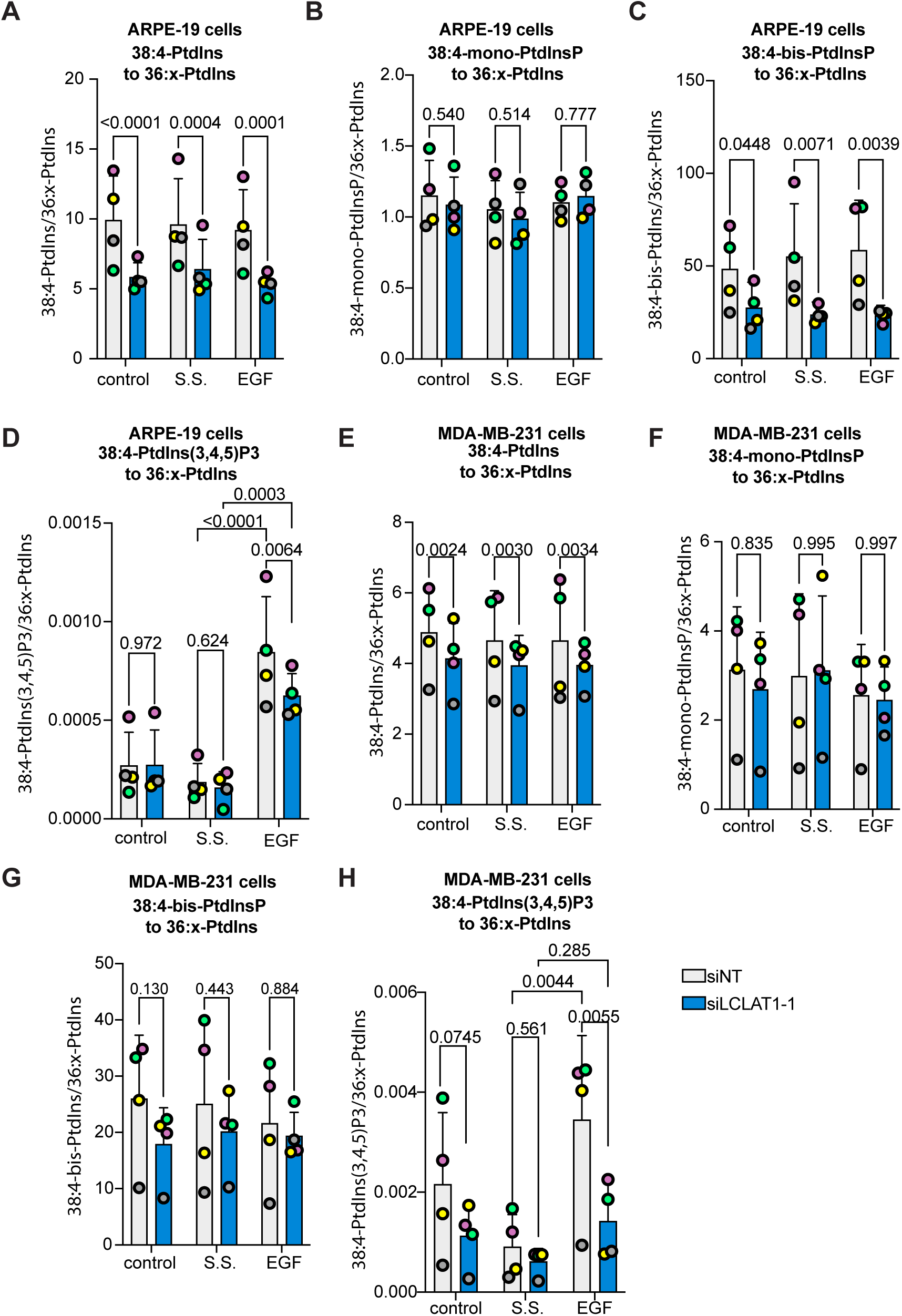
Relative levels of 38:4 PtdIns and PtdInPs in ARPE-19 and MDA-MB-231 cells silenced for LCLAT1. ARPE-19 cells (**A-D**) and MBA-MB-231 cells (**E-I**) were mock silenced (siNT) or LCLAT1-silenced. Cells were then grown in regular medium (control), serum-starved (ss), and stimulated with 5 ng/mL EGF for 5 min (EGF). Reactions were quenched and lipid extracted after addition of internal standards to primary cell extracts. PtdInsPs were measured by mass spectrometry (HPLC-MS). Shown is the ratio of standardized 38:4-PtdIns (A, E), 38:4-mono-PtdInsP (B, G), 38:4-bis-PtdInsP (C, H), and 38:4-PtdIns(3,4,5)P_3_ (D, I) to the standardized sum of 36:1 and 36:2-PtdIns (referred to as 36:x-PtdIns). Lipid analysis was repeated four independent times. Data points from matching independent experiments are colour coded. Shown are the mean ±STD. A repeated-measures, two-way ANOVA and Sidak’s post-hoc (A-C, and E-G) and Tukey’s post-hoc (D, H) tests were used to assess the data. p values are disclosed.

We similarly investigated the lipidomic profile of MDA-MB-231 cells. The major acyl species for all PtdInsPs and PtdIns was again 38:4 and we once again only detected 38:4-PtdIns(3,4,5)P_3_ acyl species in our samples. LCLAT1 suppression lowered 38:4-PtdInsP relative to 36:x-PtdInsP in MDA-MB-231 cells, but unlike ARPE-19 cells, this change did not transfer to the *bis*-species (Fig. 4G and Sup. Fig. S3D). Instead, resting cells had a significant difference in 38:4-*mono*-PtdInsPs relative to 36:x-*mono*-PtdInsPs (Sup. Fig. S3C), but not serum-starved or EGF or relative to 36:x-PtdIns (Fig. 4F and Sup. Fig. S3C). Most striking, was the elevation in the ratio of 38:4-PtdIns(3,4,5)P_3_ to 36:x-PtdInsP in non-silenced MDA-MB-231 cells after EGF stimulation and relative to serum-starved cells (Fig. 4H). Importantly, LCLAT1-perturbed MDA-MB-231 cells failed to significantly increase 38:4-PtdIns(3,4,5)P_3_ ratio compared to 36:x-PtdIns after EGF stimulation (Fig. 4H), suggesting that MDA-MB-231 cells were more sensitive to this than ARPE-19 cells. Overall, we propose that LCLAT1 expression is essential to support EGFR-dependent activation of PI3K signalling in at least two distinct cell lines, but the impact on acyl profile can vary between cell type, PtdInsP species, and treatments.

### Akt activation by EGF is defective in LCLAT1-silenced cells

Since LCLAT1-silenced cells had lower EGF-induced PtdInsP(3,4,5)P_3_ levels relative to non-silenced counterparts, we next examined if Akt activation was also impaired in ARPE-19 and MDA-MB-231 cells after EGF exposure. To do this, we probed for phosphorylation at S473 using an antibody that recognizes all isoforms of Akt when phosphorylated (pan-phospho-Akt antibody). Relative to serum-starved ARPE-19 and MDA-MB-231 cells, EGF caused a large increase in phospho-Akt in non-silenced control cells (Fig. 5A, B and Fig. 6A, B). Importantly, both ARPE-19 and MDA-MB-231 cells silenced for LCLAT1 displayed substantially reduced phospho-Akt levels after EGF stimulation (Fig. 5A, B and Fig. 6A, B). We also observed impaired p-Akt levels in ARPE-19 cells (Supplemental Figure S1A, C and Fig. 5G) and MDA-MB-231 cells (Supplemental Figure S2A, C) treated with independent oligonucleotides against LCLAT1. Since Akt1 has been reported to respond to PtdIns(3,4,5)P_3_ while Akt2 to PtdIns(3,4)P_2_ generated by SHIP2 from PtdIns(3,4,5)P_3_ (Liu et al., 2018), we sought to determine if silencing LCLAT1 exhibited any isoform specific effects on Akt phosphorylation that may reveal additional insights into PtdInsP perturbation in LCLAT1-silenced cells. We thus probed with anti-p-Akt1 (S473) and p-Akt2 (S474) antibodies to test this. We reveal that ARPE-19 and MDA-MB-231 silenced for LCLAT1 after EGF stimulation have lower levels of both p-Akt1 and p-Akt2 relative to their respective total Akt1 and Akt2 (Fig. 5C-F and Fig. 6C-F).

**Figure 5:**
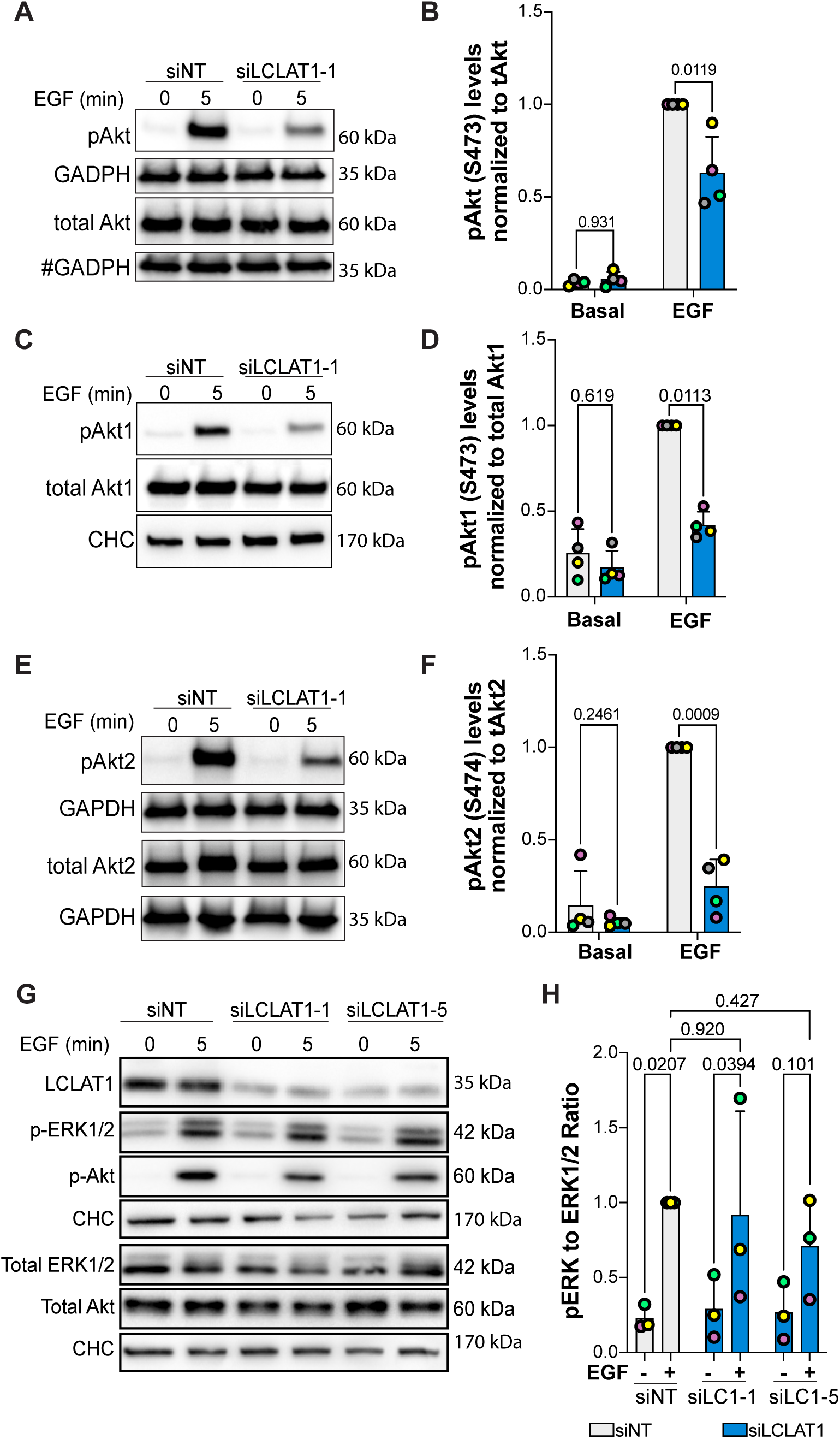
LCLAT1 is required for EGF-stimulated Akt activation in ARPE-19 cells. **A, C, E:** Mock-silenced (siNT) and LCLAT1-silenced ARPE-19 cells were serum-starved (0 min) or stimulated with 5 ng/mL EGF for 5 min. Lysates were then prepared, separated by SDS-PAGE and probed by Western blotting for pan-phospho-Akt and total pan-Akt (A), phospho-Akt1 and total Akt1 (C), and phospho-Akt2 and total Akt2 (E). Clathrin heavy chain (CHC) or GAPDH were used as loading controls. # indicates that the GAPDH blot was also used as loading control for p-TSC2 in Fig. 7G since they originated from the same membrane cut across to probe for different sized proteins. **B, D, F**: Quantification of pan-pAkt (B), pAkt1 (D), and pAkt2 (F) normalized to respective total pan-Akt, Akt1, and Akt2. **G**. ARPE-19 cells transfected with non-targeting, LCLAT1-1, or LCLAT1-5 oligonucleotides and stimulated as above. Lysates were probed with LCLAT1, phospho-ERK1/2, ERK1/2, phospho-Akt, Akt, and CHC as loading control for each blot. **E.** Quantification of phospho-ERK1/2 relative to total ERK1/2. Mean±STD are shown from n=4 (A, C, and E) and n=3 (G) independent experiments are shown. Data points from matching independent experiments are colour coded. Repeated measures two-way ANOVA and Sidak’s (B, D, F) or Tukey’s (H) post-hoc tests were used to statistically test the data. p values are indicated.

**Figure 6:**
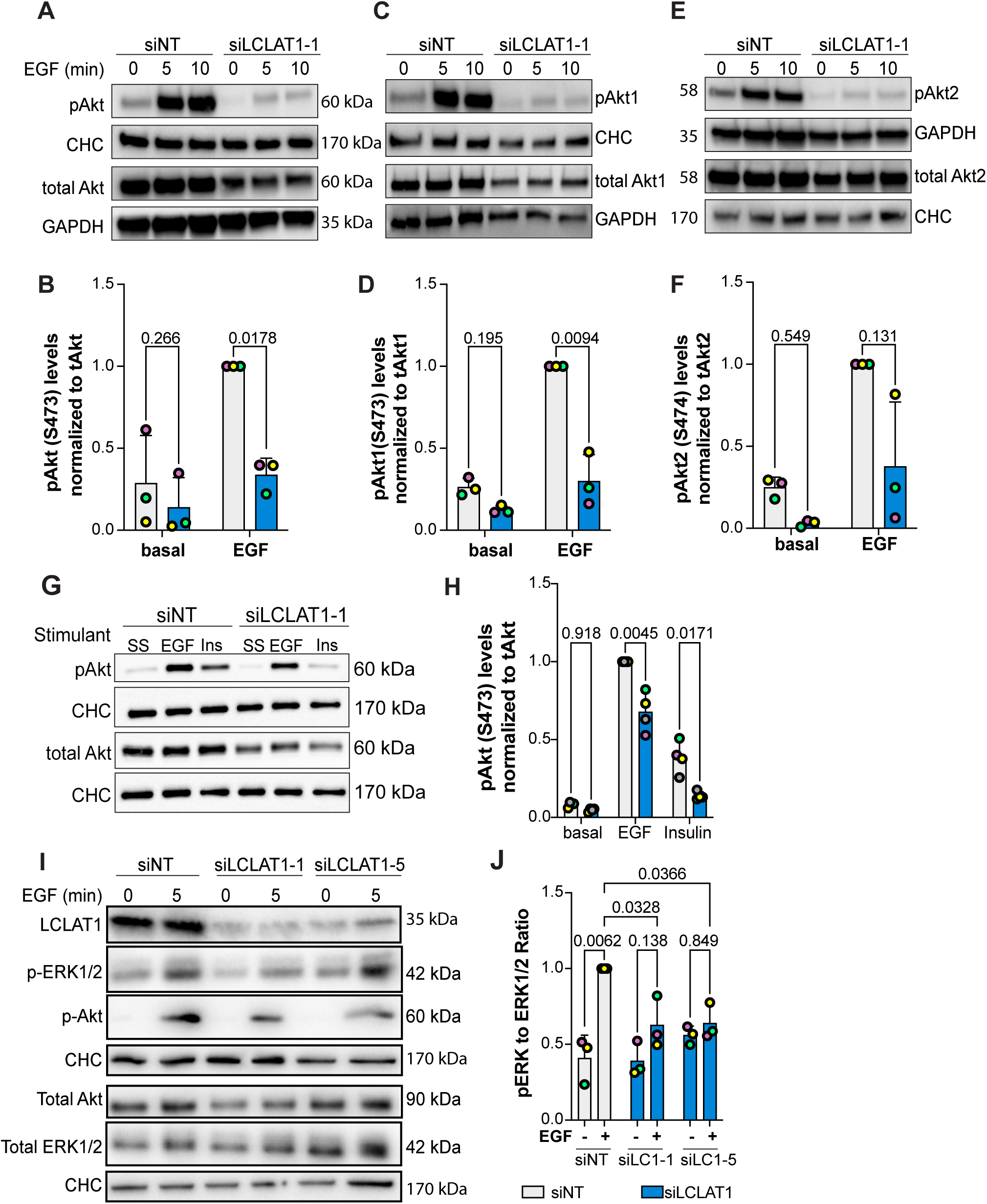
LCLAT1 is required for EGF-stimulated Akt activation in MDA-MB-231 cells. **A, C, E:** Mock-silenced and LCLAT1-silenced MDA-MB-231 cells were serum-starved (0 min) or stimulated with 5 ng/mL EGF for 5 min or 10 min. Lysates were then prepared, separated by SDS-PAGE and probed by Western blotting for pan-phospho-Akt and total pan-Akt (A), phospho-Akt1 and total Akt1 (C), and phospho-Akt2 and total Akt2 (E). Clathrin heavy chain (CHC) or GAPDH were used as loading controls. **B, D, F**: Quantification of pan-pAkt (B), pAkt1 (D), and pAkt2 (F) normalized to respective total pan-Akt, Akt1, and Akt2. **G.** Western blotting of non-silenced and LCLAT1-silenced cells after serum-starvation (SS), 5 ng/mL EGF, or 10 ng/mL insulin (Ins) stimulation for 5 min. Lysates were probed for pAkt, total Akt, and clathrin heavy chain. **E.** Quantification of pAkt relative to total Akt in treatments described in G. **I**. Western blot of MDA-MB-231 cells silenced for LCLAT1 with either LCLAT1-1 or LCLAT1-5 oligonucleotides. Cells were serum-starved or stimulated with 5 ng/mL EGF for 5 min. Lysates were probed with LCLAT1, phospho-ERK1/2, ERK1/2, phosphor-Akt, Akt, and corresponding CHC as loading control for each blot. **J**. Quantification of phospho-ERK to total ERK. Shown are the mean ±STD from n=3-4 independent experiments. Data points from matching independent experiments are colour coded. Repeated measures two-way ANOVA and Sidak’s (B, D, F) or Tukey’s (H, J) post-hoc tests were used to statistically test the data. p values are displayed.

To evince if LCLAT1 was important for Akt signaling by other receptors, we assessed insulin-mediated activation of Akt in MDA-MB-231 cells. As with EGF, LCLAT1 silencing hindered phosphorylation of Akt after insulin activation (Fig. 6G, 6H). Thus, LCLAT1 silencing negatively impacts Akt activation by both EGF and insulin signaling, implying that LCLAT1 may broadly support activation of PI3K-Akt signaling by receptor tyrosine kinases. Finally, we tested if LCLAT1-silencing also perturbed the ERK pathway by EGFR. Here, we saw cell-type specific effects. In ARPE-19 cells, EGF stimulation of phospho-ERK1/2 was predominantly unperturbed by LCLAT1 suppression with two oligonucleotides (Fig. 5G, 5H). However, in MDA-MB-231 cells inhibited for LCLAT1 displayed suppression of ERK phosphorylation in response to EGF (Fig. 6E, 6F). Overall, LCLAT1 is required for Akt activation by receptor tyrosine kinases and may play a role in ERK stimulation in a context-dependent manner.

### LCLAT1 acyltransferase silencing impairs Akt-mediated regulation of downstream targets

Since Akt activation is defective in LCLAT1-silenced cells after addition of EGF, we next examined if this effect percolated to several known Akt targets. To test for specific targets, we measured the phosphorylation state of Tsc2 and GSK3β by Western blotting. In ARPE-19 and MDA-MB-231 cells transfected with a non-targeting oligonucleotide (control siRNA), EGF promoted robust phosphorylation of these specific Akt substrates (Fig. 7). In contrast, LCLAT1 silencing led to a considerable decline in the EGF-induced phosphorylation of Tsc2 in ARPE-19 (Fig. 7A,B) and MDA-MB-231 (Fig. 7E, F) cells. For GSK3β, the effects were less clear – for ARPE-19 cells, there was a tendency for less phosphorylation of GSK3β (Fig. 7C, D), but this was not the case for MDA-MB-231 cells (Fig. 7G, H); neither cell line demonstrated a significant decrease in EGF-stimulated GSK3β phosphorylation upon LCLAT1 silencing. Overall, loss of LCLAT1 appears to compromise TSC2 regulation by Akt.

**Figure 7:**
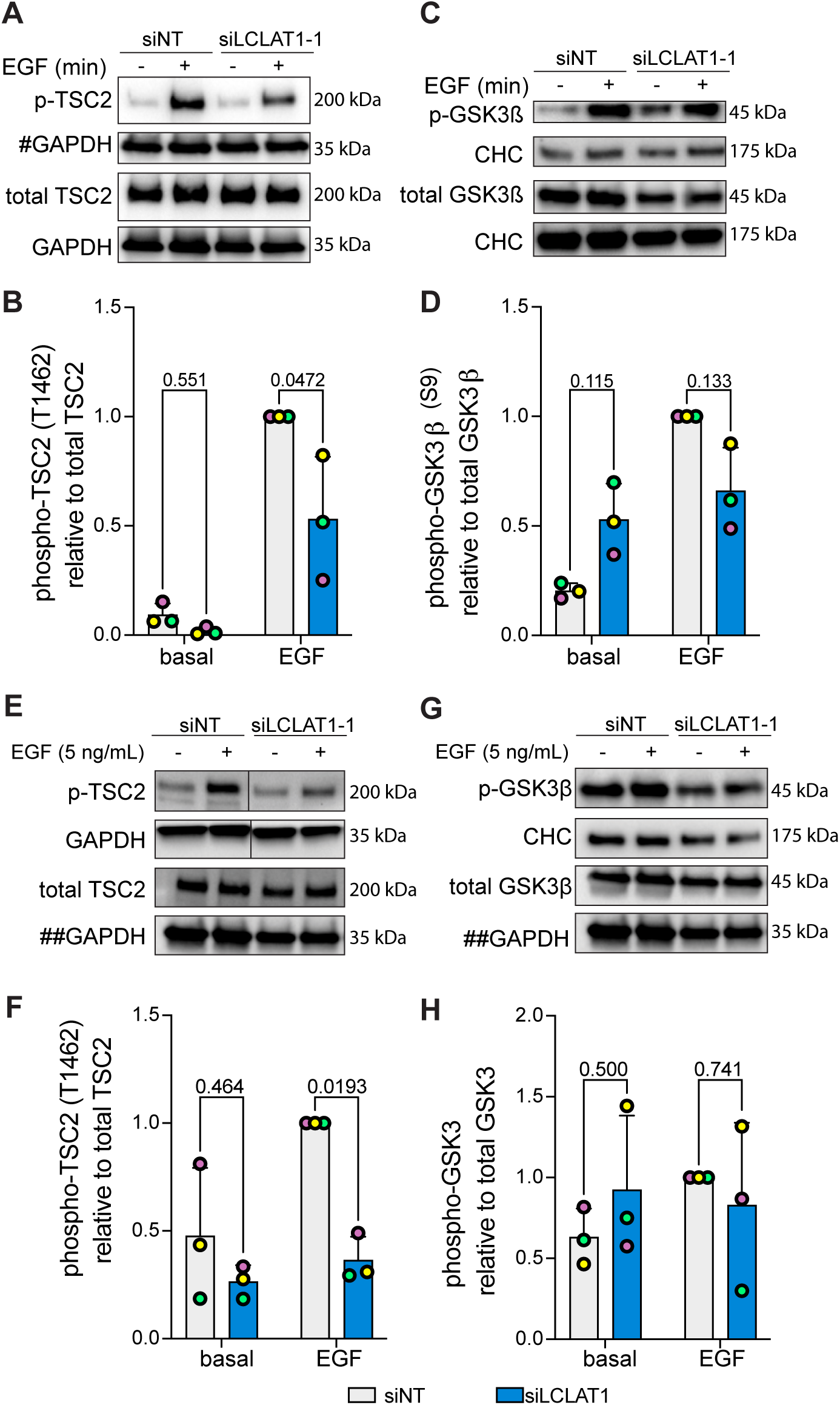
LCLAT1 is required for activation of Akt substrates after EGF stimulation. **A, C:** Mock-silenced and LCLAT1-silenced ARPE-19 cells were serum-starved (0 min) or stimulated with 5 ng/mL EGF for 5 min. Lysates were then separated by SDS-PAGE and probed by Western blotting for phospho-Tsc2 and total Tsc2 (A) and phospho-GSK3β and total GSK3β (C). Clathrin heavy chain (CHC) or GAPDH were used as loading controls. # indicates that the GAPDH blot was also used as loading control for total pan-Akt in Fig. 5A since they originated from the same membrane cut across to probe for different sized proteins. **B, D**: Quantification of pTsc2 (B) and pGSK3β (D) normalized to respective total Tsc2 and GSK3β. **E, G**: Mock-silenced and LCLAT1-silenced MDA-MB-231 cells were serum-starved (0 min) or stimulated with 5 ng/mL EGF for 5 min. Lysates were then separated by SDS-PAGE and probed by Western blotting for phospho-Tsc2 and total Tsc2 (F), and phospho-GSK3β and total GSK3β (G). Clathrin heavy chain (CHC) or GAPDH were used as loading controls. ## indicates that the GAPDH blot was used as loading control for both total TSC2 (G) and total GSK3β (I) since they originated from the same membrane cut across to probe for different sized proteins. **H, J**: Quantification of pTsc2 (H) and pGSK3β (I) normalized to respective total Tsc2 and GSK3β. For B, D, F, and H mean ± STD are shown from n=3 independent experiments. Data points from matching independent experiments are colour coded. A two-way ANOVA and Sidak’s post-hoc test was used to statistically test the data, with p values shown.

We also probed for the expression levels of cell cycle checkpoint proteins like Mdm2, p53, and p21 (Gordon et al., 2018). In doing so, we observed that LCLAT1-1 oligonucleotide, but not LCLAT1-5, tended to deplete Mdm2 protein levels in both ARPE-19 (Sup. Fig. S4A, S4B) and MDA-MB-231 cells (Supplemental Fig. S4E, S4F), despite similar levels of silencing of LCLAT1 by each LCLAT1 oligonucleotide. These oligonucleotides were significantly different from each other in their effect on p21 levels in ARPE-19 cells, with LCLAT1-1 trending upward (Supplemental Fig. S4A, S4C). There was no significant difference between oligonucleotides and non-targeting in their effect on p21 levels in MDA-MB-231 cells and for p53 protein levels in both cells (Supplemental Fig. S4). Thus, given these differences in outcome of LCLAT1 oligonucleotides that may reflect kinetics of silencing or even some limited off-target effects by the LCLAT1-1 siRNA sequence, we advise caution when considering the LCLAT1-1 oligonucleotide in future experiments. Regardless, we emphasize that the effects on Akt signaling were consistent across all oligonucleotides and in both cell types.

## Discussion

Here, we reveal that the LCLAT1 acyltransferase is needed to promote EGFR-mediated PtdIns(3,4,5)P_3_-Akt signaling in at least two cell lines. While EGFR levels and its early activation by EGF were unaffected, cells perturbed for LCLAT1 had reduced PtdIns(3,4,5)P_3_ levels and diminished Akt activation. We thus identified an important role for the poorly studied LCLAT1 acyltransferase and highlights this enzyme as part of a novel druggable lipid acyl profile remodelling pathway to modulate PI3K signalling.

### The role of LCLAT1 in generating PtdIns(3,4,5)P_3_

We provide at least three pieces of evidence that LCLAT1 is important to generate PtdIns(3,4,5)P_3_ during EGF-mediated signalling. First, we revealed that LCLAT1-silenced MDA-MB-231 and ARPE-19 cells were impaired for the recruitment of the Akt-PH-GFP, a biosensor for PtdIns(3,4,5)P_3_ levels (Haugh et al., 2000; Marshall et al., 2001), to the plasma membrane after EGF stimulation. Second, we observed that LCLAT1-silenced cells possessed lower relative levels of 38:4-PtdIns(3,4,5)P_3_. Incidentally, 38:4-PtdIns(3,4,5)P_3_ was the only observable acyl-based species for PtdIns(3,4,5)P_3_ in both cell types, likely because the other acyl isoforms of PtdIns(3,4,5)P_3_ were below the detection limit of our current analysis. Regardless, this is consistent with other studies examining the acyl profile of PtdIns(3,4,5)P_3_, identifying 38:4 as the major species, though this can vary with tissue, cell type, and genetics (Clark et al., 2011; Koizumi et al., 2019; Morioka et al., 2022; Mujalli et al., 2018). Third, Akt activation was impaired in LCLAT1-silenced cells after EGF stimulation. Hence, collectively these varied approaches indicate that LCLAT1-silenced cells are defective in promoting PtdIns(3,4,5)P_3_ levels, and possibly PtdIns(3,4)P_2_.

A key question then is how does LCLAT1 contribute to PtdIns(3,4,5)P_3_ synthesis. We propose at least three possible, non-mutually exclusive mechanisms that will need to be defined in future studies. First, enzymes involved in PtdIns(3,4,5)P_3_ metabolism such as Class I PI3Ks, the PTEN 3-phosphatase, and the SHIP1/2 5-phosphatase may display acyl sensitivity towards their substrates as previously suggested (Anderson et al., 2016), thus affecting PtdIns(3,4,5)P_3_ generation or its turnover. This was observed for Type I PIPKs, Vps34 Class III PI3Ks, and Type II phosphatases (Ohashi et al., 2020; Schmid et al., 2004; Shulga et al., 2012). Second, PtdIns(3,4,5)P_3_ levels may be reduced in LCLAT1-disturbed cells due to lower PtdIns(4,5)P_2_ substrate levels or availability, which we observed here for MDA-MB-231 cells and previously in ARPE-19 (Bone et al., 2017). This would also be consistent with Willis *et al*. who found that PtdIns(3,4,5)P_3_ signaling scales linearly with PtdIns(4,5)P_2_ levels during EGF stimulation (Wills et al., 2023). Third, and perhaps linked to the second model above, PtdIns(3,4,5)P_3_ synthesis may depend on specific substrate pools. These pools may be highly localized or even channeled through scaffolds (Choi et al., 2016) or transferred at membrane contact sites (Zaman et al., 2020). For example, reduction in PtdIns(4,5)P_2_ may lead to impaired clathrin-coated scaffold formation, which promotes EGF-mediated PI3K signaling (Cabral-Dias et al., 2022; Delos Santos et al., 2017). Indeed, we previously showed that LCLAT1 silencing altered clathrin-coated pit dynamics (Bone et al., 2017). Alternatively, contact sites between the endoplasmic reticulum and the plasma membrane are important to generate PtdIns(4,5)P_2_ (Chang and Liou, 2015; Cockcroft et al., 2016; Kim et al., 2015; Lees et al., 2017; Saheki et al., 2016; Zaman et al., 2020). These sites may provide precursor pools for PtdIns(4,5)P_2_ and/or PtdIns(3,4,5)P_3_; consistent with this, depletion of bulk PtdIns(4)P from the plasma membrane did not reduce PtdIns(4,5)P_2_ levels (Hammond et al., 2012), intimating that PtdIns(4,5)P_2_ depends on specific pools of PtdIns(4)P. Consistent with this, we previously noted colocalization of a subset of LCLAT1 with proteins known to be at endoplasmic reticulum-plasma membrane contact sites, such as extended synaptotagmins (E-Syt2), (Bone et al., 2017). Thus, LCLAT1 may play a role in generating specific substrate pools to support PtdIns(4,5)P_2_ and PtdIns(3,4,5)P_3_ at the plasma membrane.

The notion that LCLAT1 may act on specific pools of PtdInsPs is consistent with observations that LCLAT1 and LPIAT/MBOAT7 do not affect the levels of and the acyl profile of all PtdInsPs. For example, their disruption preferentially affects the acyl profile of PtdIns and/or *bis*-phosphorylated PtdInsPs, but not of mono-PtdInsPs in ARPE-19 cells (Anderson et al., 2013; Bone et al., 2017) (Fig. 4, Sup. Fig. S3). By comparison, LCLAT1 disruption altered 38:4-PtdIns and 38:4-PtdIns(3,4,5)P_3_ in MDA-MB-231 cells, but had little effect on other PtdInsPs in most conditions tested. Thus, LCLAT1 may act on specialized pools of lipids, but this is likely cell-type specific, while enzymes like CDS2 and DGKε may also play a role in establishing PtdInsP acyl profiles (Bozelli and Epand, 2019; D’Souza et al., 2014; Shulga et al., 2011). Overall, while we know that LCLAT1 is needed to boost PtdIns(3,4,5)P_3_ levels in response to EGF, the exact mechanism of action remains to be defined.

### LCLAT1 and PtdIns(3,4,5)P_3_ functions

We witnessed that LCLAT1 supports PtdIns(3,4,5)P_3_-mediated activation of Akt isoforms after EGF signalling. Consequently, TSC2 a key target of Akt was less phosphorylated in LCLAT1-silenced cells. Hence, we anticipate that LCLAT1 affects other effector functions of PtdIns(3,4,5)P_3_ including activation of other kinases such as Btk and GEFs for the Rho-family of GTPases such as Vav1 and Tiam1 (Salamon and Backer, 2013; Wang et al., 2006, 2; Zhu et al., 2015, 1). Additionally, while our work focused on EGFR-mediated signalling, we observed reduced insulin-driven activation of Akt as well (Sup. Fig. S2D, E). Thus, we postulate that LCLAT1 broadly supports PtdInsP-dependent signalling by other receptor tyrosine kinases, and may also support such signalling by GPCRs and peripherally-associated kinase receptors such as immune receptors (Bresnick and Backer, 2019; Dowling and Mansell, 2016; Getahun and Cambier, 2015; Takeuchi and Ito, 2011). Conceivably, the putative role of LCLAT1 in promoting PI3K signalling among these receptor classes may depend on which isoforms of Class I and/or Class II PI3K are engaged (Bilanges et al., 2019; Duncan et al., 2020). Overall, our observations establish a key relationship between LCLAT1 and EGFR-PtdIns(3,4,5)P_3_-Akt axis and sets a course to determine the universality of LCLAT1 acyltransferase in PtdIns(3,4,5)P_3_ signalling.

### LCLAT1 in cellular function and therapeutic potential

While the LCLAT1 acyltransferase remains relatively under-investigated, LCLAT1 is associated with a variety of functions (Zhang et al., 2023). These include hematopoiesis and cell differentiation (Huang et al., 2014; Huang et al., 2017; Wang et al., 2007; Xiong et al., 2008), metabolic regulation (Cao et al., 2009, 1; Liu et al., 2012), mitochondrial stability, dynamics and function (Huang et al., 2020; Li et al., 2010, 1; Li et al., 2012), sensitivity to oxidative stress (Li et al., 2010; Liu et al., 2012), endocytosis and endosomal trafficking (Bone et al., 2017), and now receptor tyrosine kinase signalling. LCLAT1 is also proposed to remodel the acyl profile of both cardiolipin and PtdIns/PtdInsPs (Bone et al., 2017; Cao et al., 2004, 1; Imae et al., 2012; Li et al., 2010). It is generally thought that mitochondrial and oxidative stress occurs through cardiolipin remodelling, while endocytosis and receptor signalling is connected to PtdIns acyl function, as proposed here. However, the specific roles of LCLAT1 in cardiolipin and PtdInsP acyl remodelling have not been reconciled. It may be that LCLAT1 has independent roles in acylating these two distinct lipids, or alternatively, one may depend on the other. For example, mitochondria contain PtdIns on their outer membrane, which is important for mitochondria dynamics and function (Pemberton et al., 2020; Zewe et al., 2020). Conceivably, then LCLAT1 acylation of PtdIns may impact lipidomic properties of cardiolipin in mitochondria. In addition, while we did not observe significant changes in Mdm2, p53 and p21 levels that we could specifically attribute to LCLAT1 disturbance, it will be important to examine the effect of LCLAT1 suppression in cell cycle and apoptosis. Unfortunately, we eventually discovered that LCLAT1-1 oligonucleotide appears to have either distinct kinetics of knockdown or a limited set of non-specific effects on Mdm2 protein levels. Regardless, we have performed extensive experiments that allowed us to ascertain that the effects on PI3K-Akt signaling were observed with multiple LCLAT1 silencing oligonucleotides. Overall, LCLAT1 and its putative partner, MBOAT7/LPIAT1, remain relatively understudied and without well-established inhibitors. Given the role of LCLAT1 in PtdInsP biology and receptor signalling, we propose that these acyltransferases represent new targets for therapeutic development.

## Supplemental Information

**Supplemental Figure S1: LCLAT1 silencing in ARPE-19 cells with an independent siRNA oligonucleotide disrupts Akt signalling.** ARPE-19 cells were transfected with oligonucleotide siLCLAT1-5 or non-targeting control. Cells were then serum-starved (SS), followed by 5 ng/mL EGF stimulation for 5 min. **A.** Lysates were probed for LCLAT1 expression, p-Akt, Akt, and clathrin heavy chain (CHC), which was used as a loading control. **B.** Quantification of LCLAT1 silencing by normalizing LCLAT1 to CHC signal in ARPE-19 cells. **C.** Quantification of p-Akt levels relative to total Akt. Data are shown as mean ±STD are shown from n=3 independent experiments. Data points from matching independent experiments are colour coded. Data was analysed by a repeated measures two-way ANOVA and Sidak’s post-hoc test. p values are disclosed.

**Supplemental Figure S2: LCLAT1 silencing in MDA-MB-231 cells with independent siRNA oligonucleotides disrupts Akt signalling. A.** Western blotting showing LCLAT1 silencing in MDA-MB-231 cells transfected with non-targeting siRNA, siLCLAT1-1, siLCLAT1-2, or siLCLAT1-3 oligonucleotides. Cells were then serum-starved (SS) or stimulated with 5 ng/mL EGF for 5 min. Lysates were probed for LCLAT1 expression, p-Akt, and clathrin heavy chain (CHC), which was used as a loading control. **B.** Quantification of LCLAT1 silencing by normalizing LCLAT1 to CHC signal in MDA-MB-231 cells transfected as in D. C. Quantification of p-Akt levels relative to clathrin in MDA-MB-231 cells mock-silenced or LCLAT1-silenced with one of three siRNA oligonucleotides and either serum-starved or stimulated with 5 ng/mL EGF for 5 min. Data points from matching independent experiments are colour coded. Data in B were analysed with a repeated measures one-way ANOVA and Dunnett’s post-hoc test. Data in C were analysed with a repeated measures two-way ANOVA and Tukey’s post-hoc test. p values are indicated.

**Supplemental Figure S3. Relative levels of 38:4-PtdInsPs to 36:x-PtdInsPs in ARPE-19 and MDA-MB-231 cells silenced for LCLAT1.** ARPE-19 cells (**A-B**) and MBA-MB-231 cells (**C-D**) were mock silenced (siCon) or LCLAT1-silenced. Cells were then grown in regular medium (control), serum-starved (ss), and stimulated with 5 ng/mL EGF for 5 min (EGF). Reactions were quenched and lipid extracted after addition of internal standards to primary cell extracts. PtdInsPs were measured by mass spectrometry (HPLC-MS). Shown is the ratio of standardized 38:4-mono-PtdIns (A, C) and 38:4-bis-PtdInsP2 (B, D) to the respective standardized sum of 36:1 and 36:2- (referred to as 36:x-PtdIns) mono-PtdInsP and bis-PtdInsP2. Lipid analysis was repeated four independent times. Data points from matching independent experiments are colour coded. Shown are the mean ±STD. A repeated measures two-way ANOVA and Sidak’s post-hoc test was used to test data. p values are indicated.

**Supplemental Figure S4: LCLAT1 silencing effect on cell cycle checkpoint proteins, Mdm2, p21, and p53. A, B.** Western blotting showing LCLAT1 silencing in ARPE-19 (A) and MDA-MB-231 (B) cells transfected with non-targeting siRNA, siLCLAT1-1, or siLCLAT1-5 oligonucleotides. Cells were serum-starved or stimulated with 5 ng/mL EGF for 5 min. Lysates were probed for LCLAT1, Mdm2, p53, and p21 expression. Clathrin heavy chain (CHC) was probed as a loading control. **B, C, D, F, G, H.** Quantification of Mdm2, p21, and p53 expression relative to CHC expression in ARPE-19 (B-D) and MDA-MB-231 (F-H) cells. Data are the mean ± STD from n=4 independent experiments. Data points from matching independent experiments are colour coded. Data was analysed with a repeated measures two-way ANOVA and Tukey’s post-hoc test with p values indicated.

## References

Alessi, D. R., Deak, M., Casamayor, A., Caudwell, F. B., Morrice, N., Norman, D. G., Gaffney, P., Reese, C. B., MacDougall, C. N., Harbison, D., et al. (1997). 3-Phosphoinositide-dependent protein kinase-1 (PDK1): Structural and functional homology with the Drosophila DSTPK61 kinase. Current Biology 7, 776–789.

Anderson, K. E., Kielkowska, A., Durrant, T. N., Juvin, V., Clark, J., Stephens, L. R. and Hawkins, P. T. (2013). Lysophosphatidylinositol-Acyltransferase-1 (LPIAT1) Is Required to Maintain Physiological Levels of PtdIns and PtdInsP2 in the Mouse. PLoS ONE 8, e58425.

Anderson, K. E., Juvin, V., Clark, J., Stephens, L. R. and Hawkins, P. T. (2016). Investigating the effect of arachidonate supplementation on the phosphoinositide content of MCF10a breast epithelial cells. Advances in biological regulation 62, 18–24.

Balla, T. (2013). Phosphoinositides: tiny lipids with giant impact on cell regulation. Physiological reviews 93, 1019–137.

Barneda, D., Cosulich, S., Stephens, L. and Hawkins, P. (2019). How is the acyl chain composition of phosphoinositides created and does it matter? Biochemical Society Transactions 47, 1291–1305.

Bellacosa, A., Chan, T. O., Ahmed, N. N., Datta, K., Malstrom, S., Stokoe, D., McCormick, F., Feng, J. and Tsichlis, P. (1998). Akt activation by growth factors is a multiple-step process: the role of the PH domain. Oncogene 17, 313–325.

Bilanges, B., Posor, Y. and Vanhaesebroeck, B. (2019). PI3K isoforms in cell signalling and vesicle trafficking. Nat Rev Mol Cell Biol 20, 515–534.

Blunsom, N. J. and Cockcroft, S. (2020). Phosphatidylinositol synthesis at the endoplasmic reticulum. Biochimica et Biophysica Acta - Molecular and Cell Biology of Lipids 1865,.

Bone, L. N., Dayam, R. M., Lee, M., Kono, N., Fairn, G. D., Arai, H., Botelho, R. J. and Antonescu, C. N. (2017). The acyltransferase LYCAT controls specific phosphoinositides and related membrane traffic. Molecular Biology of the Cell 28, 161–172.

Böni-Schnetzler, M. and Pilch, P. F. (1987). Mechanism of epidermal growth factor receptor autophosphorylation and high-affinity binding. Proc Natl Acad Sci U S A 84, 7832–7836.

Bozelli, J. C. and Epand, R. M. (2019). Specificity of Acyl Chain Composition of Phosphatidylinositols. Proteomics 19,.

Bresnick, A. R. and Backer, J. M. (2019). PI3Kβ-A Versatile Transducer for GPCR, RTK, and Small GTPase Signaling. Endocrinology 160, 536–555.

Cabral-Dias, R., Awadeh, Y., Botelho, R. J. and Antonescu, C. N. (2021). Detection of Plasma Membrane Phosphoinositide Dynamics Using Genetically Encoded Fluorescent Protein Probes. In Methods in Molecular Biology, pp. 73–89. Humana Press Inc.

Cabral-Dias, R., Lucarelli, S., Zak, K., Rahmani, S., Judge, G., Abousawan, J., DiGiovanni, L. F., Vural, D., Anderson, K. E., Sugiyama, M. G., et al. (2022). Fyn and TOM1L1 are recruited to clathrin-coated pits and regulate Akt signaling. Journal of Cell Biology 221,.

Cao, J., Liu, Y., Lockwood, J., Burn, P. and Shi, Y. (2004). A novel cardiolipin-remodeling pathway revealed by a gene encoding an endoplasmic reticulum-associated acyl-CoA:lysocardiolipin acyltransferase (ALCAT1) in mouse. Journal of Biological Chemistry 279, 31727–31734.

Cao, J., Shen, W., Chang, Z. and Shi, Y. (2009). ALCAT1 is a polyglycerophospholipid acyltransferase potently regulated by adenine nucleotide and thyroid status. American Journal of Physiology-Endocrinology and Metabolism 296, E647–E653.

Chang, C. L. and Liou, J. (2015). Phosphatidylinositol 4, 5-bisphosphate homeostasis regulated by Nir2 and Nir3 proteins at endoplasmic reticulum-plasma membrane junctions. Journal of Biological Chemistry 290, 14289–14301.

Choi, S., Hedman, A. C., Sayedyahossein, S., Thapa, N., Sacks, D. B. and Anderson, R. A. (2016). Agonist-stimulated phosphatidylinositol-3,4,5-trisphosphate generation by scaffolded phosphoinositide kinases. Nat Cell Biol 18, 1324–1335.

Choy, C. H., Han, B. K. and Botelho, R. J. (2017). Phosphoinositide Diversity, Distribution, and Effector Function: Stepping Out of the Box. BioEssays 39, 1700121.

Clark, J., Anderson, K. E., Juvin, V., Smith, T. S., Karpe, F., Wakelam, M. J. O., Stephens, L. R. and Hawkins, P. T. (2011). Quantification of PtdInsP3 molecular species in cells and tissues by mass spectrometry. Nature Methods 8, 267–272.

Cockcroft, S., Garner, K., Yadav, S., Gomez-Espinoza, E. and Raghu, P. (2016). RdgB reciprocally transfers PA and PI at ER-PM contact sites to maintain PI(4,5)P2 homoeostasis during phospholipase C signalling in Drosophila photoreceptors. Biochemical Society Transactions 44, 286–292.

Cross, D. A. E., Alessi, D. R., Cohen, P., Andjelkovich, M. and Hemmings, B. A. (1995). Inhibition of glycogen synthase kinase-3 by insulin mediated by protein kinase B. Nature 378, 785–789.

Delos Santos, R. C., Bautista, S., Lucarelli, S., Bone, L. N., Dayam, R. M., Abousawan, J., Botelho, R. J. and Antonescu, C. N. (2017). Selective regulation of clathrin-mediated epidermal growth factor receptor signaling and endocytosis by phospholipase C and calcium. Mol Biol Cell 28, 2802–2818.

Dey, N., De, P. and Leyland-Jones, B. (2017). PI3K-AKT-mTOR inhibitors in breast cancers: From tumor cell signaling to clinical trials. Pharmacology and Therapeutics 175, 91–106.

Dibble, C. C. and Manning, B. D. (2013). Signal integration by mTORC1 coordinates nutrient input with biosynthetic output. Nature cell biology 15, 555–64.

Dickson, E. J. and Hille, B. (2019). Understanding phosphoinositides: rare, dynamic, and essential membrane phospholipids. Biochem J 476, 1–23.

Doumane, M., Caillaud, M. C. and Jaillais, Y. (2022). Experimental manipulation of phosphoinositide lipids: from cells to organisms. Trends in Cell Biology 32, 445–461.

Dowling, J. K. and Mansell, A. (2016). Toll-like receptors: the swiss army knife of immunity and vaccine development. Clinical & Translational Immunology 5, e85.

D’Souza, K. and Epand, R. M. (2014). Enrichment of phosphatidylinositols with specific acyl chains. Biochim. Biophys. Acta - Biomembr. 1838, 1501–1508.

D’Souza, K., Kim, Y. J., Balla, T. and Epand, R. M. (2014). Distinct properties of the two isoforms of CDP-diacylglycerol synthase. Biochemistry 53, 7358–7367.

Duncan, L., Shay, C. and Teng, Y. (2020). PI3K Isoform-Selective Inhibitors in Cancer. Adv Exp Med Biol 1255, 165–173.

Freyr Eiriksson, F., Kampp Nøhr, M., Costa, M., Klara Bö dvarsdottir, S., Margret, H. O. and Thorsteinsdottir, M. (2020). Lipidomic study of cell lines reveals differences between breast cancer subtypes.

Getahun, A. and Cambier, J. C. (2015). Of ITIMs, ITAMs, and ITAMis: Revisiting immunoglobulin Fc receptor signaling. Immunological Reviews 268, 66–73.

Gordon, E. M., Ravicz, J. R., Liu, S., Chawla, S. P. and Hall, F. L. (2018). Cell cycle checkpoint control: The cyclin G1/Mdm2/p53 axis emerges as a strategic target for broad-spectrum cancer gene therapy - A review of molecular mechanisms for oncologists. Mol Clin Oncol 9, 115–134.

Gullick, W. J., Downward, J. and Waterfield, M. D. (1985). Antibodies to the autophosphorylation sites of the epidermal growth factor receptor protein-tyrosine kinase as probes of structure and function. EMBO J 4, 2869–2877.

Haag, M., Schmidt, A., Sachsenheimer, T. and Brügger, B. (2012). Quantification of Signaling Lipids by Nano-Electrospray Ionization Tandem Mass Spectrometry (Nano-ESI MS/MS). Metabolites 2, 57–76.

Hammond, G. R. V., Fischer, M. J., Anderson, K. E., Holdich, J., Koteci, A., Balla, T. and Irvine, R. F. (2012). PI4P and PI(4,5)P2 are essential but independent lipid determinants of membrane identity. *Science (New York*, N.Y.) 337, 727–30.

Haugh, J. M., Codazzi, F., Teruel, M. and Meyer, T. (2000). Spatial Sensing in Fibroblasts Mediated by 3′ Phosphoinositides. Journal of Cell Biology 151, 1269–1280.

Hicks, A. M., DeLong, C. J., Thomas, M. J., Samuel, M. and Cui, Z. (2006). Unique molecular signatures of glycerophospholipid species in different rat tissues analyzed by tandem mass spectrometry. Biochimica et Biophysica Acta (BBA) - Molecular and Cell Biology of Lipids 1761, 1022–1029.

Holgado-Madruga, M., Emlet, D. R., Moscatello, D. K., Godwin, A. K. and Wong, A. J. (1996). A Grb2-associated docking protein in EGF- and insulin-receptor signalling. Nature 379, 560–564.

Honegger, A. M., Szapary, D., Schmidt, A., Lyall, R., Van Obberghen, E., Dull, T. J., Ullrich, A. and Schlessinger, J. (1987). A mutant epidermal growth factor receptor with defective protein tyrosine kinase is unable to stimulate proto-oncogene expression and DNA synthesis. Mol Cell Biol 7, 4568–4571.

Hu, P., Margolis, B., Skolnik, E. Y., Lammers, R., Ullrich, A. and Schlessinger, J. (1992). Interaction of phosphatidylinositol 3-kinase-associated p85 with epidermal growth factor and platelet-derived growth factor receptors. Mol Cell Biol 12, 981–990.

Huang, L. S., Mathew, B., Li, H., Zhao, Y., Ma, S. F., Noth, I., Reddy, S. P., Harijith, A., Usatyuk, P. V., Berdyshev, E. V., et al. (2014). The mitochondrial cardiolipin remodeling enzyme lysocardiolipin acyltransferase is a novel target in pulmonary fibrosis. American Journal of Respiratory and Critical Care Medicine 189, 1402–1415.

Huang, L. S., Jiang, P., Feghali-Bostwick, C., Reddy, S. P., Garcia, J. G. N. and Natarajan, V. (2017). Lysocardiolipin acyltransferase regulates TGF-β mediated lung fibroblast differentiation. Free Radical Biology and Medicine 112, 162–173.

Huang, L. S., Kotha, S. R., Avasarala, S., VanScoyk, M., Winn, R. A., Pennathur, A., Yashaswini, P. S., Bandela, M., Salgia, R., Tyurina, Y. Y., et al. (2020). Lysocardiolipin acyltransferase regulates NSCLC cell proliferation and migration by modulating mitochondrial dynamics. Journal of Biological Chemistry 295, 13393–13406.

Idevall-Hagren, O. and De Camilli, P. (2015). Detection and manipulation of phosphoinositides. Biochimica et Biophysica Acta - Molecular and Cell Biology of Lipids 1851, 736–745.

Imae, R., Inoue, T., Nakasaki, Y., Uchida, Y., Ohba, Y., Kono, N., Nakanishi, H., Sasaki, T., Mitani, S. and Arai, H. (2012). LYCAT, a homologue of *C. elegans acl-8*, *acl-9*, and *acl-10*, determines the fatty acid composition of phosphatidylinositol in mice. Journal of Lipid Research 53, 335–347.

Inoki, K., Li, Y., Zhu, T., Wu, J. and Guan, K.-L. (2002). TSC2 is phosphorylated and inhibited by Akt and suppresses mTOR signalling. Nature Cell Biology 4, 648–657.

Inoki, K., Li, Y., Xu, T. and Guan, K. L. (2003). Rheb GTpase is a direct target of TSC2 GAP activity and regulates mTOR signaling. Genes and Development 17, 1829–1834.

Katan, M. and Cockcroft, S. (2020). Phosphatidylinositol(4,5)bisphosphate: diverse functions at the plasma membrane. Essays in Biochemistry 20200041.

Kim, Y. J., Guzman-Hernandez, M.-L., Wisniewski, E. and Balla, T. (2015). Phosphatidylinositol-Phosphatidic Acid Exchange by Nir2 at ER-PM Contact Sites Maintains Phosphoinositide Signaling Competence. Developmental Cell 33, 549–561.

Kiyatkin, A., Aksamitiene, E., Markevich, N. I., Borisov, N. M., Hoek, J. B. and Kholodenko, B. N. (2006). Scaffolding protein Grb2-associated binder 1 sustains epidermal growth factor-induced mitogenic and survival signaling by multiple positive feedback loops. Journal of Biological Chemistry 281, 19925–19938.

Koizumi, A., Narita, S., Nakanishi, H., Ishikawa, M., Eguchi, S., Kimura, H., Takasuga, S., Huang, M., Inoue, T., Sasaki, J., et al. (2019). Increased fatty acyl saturation of phosphatidylinositol phosphates in prostate cancer progression. Scientific Reports 9, 1– 8.

Koland, J. G. and Cerione, R. A. (1988). Growth factor control of epidermal growth factor receptor kinase activity via an intramolecular mechanism. J Biol Chem 263, 2230–2237.

Lee, H.-C., Inoue, T., Sasaki, J., Kubo, T., Matsuda, S., Nakasaki, Y., Hattori, M., Tanaka, F., Udagawa, O., Kono, N., et al. (2012). LPIAT1 regulates arachidonic acid content in phosphatidylinositol and is required for cortical lamination in mice. Molecular biology of the cell 23, 4689–700.

Lees, J. A., Messa, M., Sun, E. W., Wheeler, H., Torta, F., Wenk, M. R., De Camilli, P. and Reinisch, K. M. (2017). Lipid transport by TMEM24 at ER-plasma membrane contacts regulates pulsatile insulin secretion. Science 355, eaah6171.

Li, J., Romestaing, C., Han, X., Li, Y., Hao, X., Wu, Y., Sun, C., Liu, X., Jefferson, L. S., Xiong, J., et al. (2010). Cardiolipin remodeling by ALCAT1 links oxidative stress and mitochondrial dysfunction to obesity. Cell Metabolism 12, 154–165.

Li, J., Liu, X., Wang, H., Zhang, W., Chan, D. C. and Shi, Y. (2012). Lysocardiolipin acyltransferase 1 (ALCAT1) controls mitochondrial DNA fidelity and biogenesis through modulation of MFN2 expression. Proceedings of the National Academy of Sciences of the United States of America 109, 6975–6980.

Li, S., Shen, Y., Wang, M., Yang, J., Lv, M., Li, P., Chen, Z. and Yang, J. (2017). Loss of PTEN expression in breast cancer: association with clinicopathological characteristics and prognosis. Oncotarget 8, 32043–32054.

Linggi, B. and Carpenter, G. (2006). ErbB receptors: new insights on mechanisms and biology. Trends in Cell Biology 16, 649–656.

Liu, X., Ye, B., Miller, S., Yuan, H., Zhang, H., Tian, L., Nie, J., Imae, R., Arai, H., Li, Y., et al. (2012). Ablation of ALCAT1 Mitigates Hypertrophic Cardiomyopathy through Effects on Oxidative Stress and Mitophagy. Molecular and Cellular Biology 32, 4493–4504.

Liu, S. L., Wang, Z. G., Hu, Y., Xin, Y., Singaram, I., Gorai, S., Zhou, X., Shim, Y., Min, J. H., Gong, L. W., et al. (2018). Quantitative Lipid Imaging Reveals a New Signaling Function of Phosphatidylinositol-3,4-Bisphophate: Isoform- and Site-Specific Activation of Akt. Molecular Cell 71, 1092–1104.e5.

Manning, B. D. and Toker, A. (2017). AKT/PKB Signaling: Navigating the Network. Cell 169, 381–405.

Margolis, B., Bellot, F., Honegger, A. M., Ullrich, A., Schlessinger, J. and Zilberstein, A. (1990a). Tyrosine kinase activity is essential for the association of phospholipase C-gamma with the epidermal growth factor receptor. Mol Cell Biol 10, 435–441.

Margolis, B., Li, N., Koch, A., Mohammadi, M., Hurwitz, D. R., Zilberstein, A., Ullrich, A., Pawson, T. and Schlessinger, J. (1990b). The tyrosine phosphorylated carboxyterminus of the EGF receptor is a binding site for GAP and PLC-gamma. EMBO J 9, 4375–4380.

Marshall, J. G., Booth, J. W., Stambolic, V., Mak, T., Balla, T., Schreiber, A. D., Meyer, T. and Grinstein, S. (2001). Restricted accumulation of phosphatidylinositol 3-kinase products in a plasmalemmal subdomain during Fc?? receptor-mediated phagocytosis. Journal of Cell Biology 153, 1369–1380.

Milne, S. B., Ivanova, P. T., DeCamp, D., Hsueh, R. C. and Brown, H. A. (2005). A targeted mass spectrometric analysis of phosphatidylinositol phosphate species. J Lipid Res 46, 1796–1802.

Morioka, S., Nakanishi, H., Yamamoto, T., Hasegawa, J., Tokuda, E., Hikita, T., Sakihara, T., Kugii, Y., Oneyama, C., Yamazaki, M., et al. (2022). A mass spectrometric method for in-depth profiling of phosphoinositide regioisomers and their disease-associated regulation. Nat Commun 13, 83.

Mujalli, A., Chicanne, G., Bertrand-Michel, J., Viars, F., Stephens, L., Hawkins, P., Viaud, J., Gaits-Iacovoni, F., Severin, S., Gratacap, M. P., et al. (2018). Profiling of phosphoinositide molecular species in human and mouse platelets identifies new species increasing following stimulation. Biochimica et Biophysica Acta - Molecular and Cell Biology of Lipids 1863, 1121–1131.

Naguib, A., Bencze, G., Engle, D. D., Chio, I. I. C., Herzka, T., Watrud, K., Bencze, S., Tuveson, D. A., Pappin, D. J. and Trotman, L. C. (2015). P53 mutations change phosphatidylinositol acyl chain composition. Cell Reports 10, 8–19.

Ohashi, Y., Tremel, S., Masson, G. R., McGinney, L., Boulanger, J., Rostislavleva, K., Johnson, C. M., Niewczas, I., Clark, J. and Williams, R. L. (2020). Membrane characteristics tune activities of endosomal and autophagic human VPS34 complexes. eLife 9, e58281.

Orofiamma, L. A., Vural, D. and Antonescu, C. N. (2022). Control of cell metabolism by the epidermal growth factor receptor. Biochim Biophys Acta Mol Cell Res 119359.

Pemberton, J. G., Kim, Y. J., Humpolickova, J., Eisenreichova, A., Sengupta, N., Toth, D. J., Boura, E. and Balla, T. (2020). Defining the subcellular distribution and metabolic channeling of phosphatidylinositol. Journal of Cell Biology 219,.

Posor, Y., Jang, W. and Haucke, V. (2022). Phosphoinositides as membrane organizers. Nat Rev Mol Cell Biol.

Rodgers, S. J., Ferguson, D. T., Mitchell, C. A. and Ooms, L. M. (2017). Regulation of PI3K effector signalling in cancer by the phosphoinositide phosphatases. Bioscience Reports 37, BSR20160432.

Rodrigues, G. A., Falasca, M., Zhang, Z., Ong, S. H. and Schlessinger, J. (2000). A novel positive feedback loop mediated by the docking protein Gab1 and phosphatidylinositol 3-kinase in epidermal growth factor receptor signaling. Mol Cell Biol 20, 1448–1459.

Rueda-Rincon, N., Bloch, K., Derua, R., Vyas, R., Harms, A., Hankemeier, T., Ali Khan, N., Dehairs, J., Bagadi, M., Mercedes Binda, M., et al. (2015). p53 attenuates AKT signaling by modulating membrane phospholipid composition. Oncotarget 6, 21240–21254.

Saheki, Y., Bian, X., Schauder, C. M., Sawaki, Y., Surma, M. A., Klose, C., Pincet, F., Reinisch, K. M. and De Camilli, P. (2016). Control of plasma membrane lipid homeostasis by the extended synaptotagmins. Nature Cell Biology 18, 504–515.

Salamon, R. S. and Backer, J. M. (2013). Phosphatidylinositol-3,4,5-trisphosphate: Tool of choice for class I PI 3-kinases. BioEssaysJ: news and reviews in molecular, cellular and developmental biology 35, 602–11.

Schindelin, J., Arganda-Carreras, I., Frise, E., Kaynig, V., Longair, M., Pietzsch, T., Preibisch, S., Rueden, C., Saalfeld, S., Schmid, B., et al. (2012). Fiji: An open-source platform for biological-image analysis. Nature Methods 9, 676–682.

Schmid, A. C., Wise, H. M., Mitchell, C. A., Nussbaum, R. and Woscholski, R. (2004). Type II phosphoinositide 5-phosphatases have unique sensitivities towards fatty acid composition and head group phosphorylation. FEBS Letters 576, 9–13.

Shulga, Y. V., Topham, M. K. and Epand, R. M. (2011). Substrate specificity of diacylglycerol kinase-epsilon and the phosphatidylinositol cycle. FEBS Lett 585, 4025–4028.

Shulga, Y. V., Anderson, R. A., Topham, M. K. and Epand, R. M. (2012). Phosphatidylinositol-4-phosphate 5-kinase isoforms exhibit acyl chain selectivity for both substrate and lipid activator. J. Biol. Chem. 287, 35953–35963.

Stauffer, T. P., Ahn, S. and Meyer, T. (1998). Receptor-induced transient reduction in plasma membrane PtdIns(4,5)P2 concentration monitored in living cells. Curr. Biol. 8, 343–346.

Stokoe, D., Stephens, L. R., Copeland, T., Gaffney, P. R. J., Reese, C. B., Painter, G. F., Holmes, A. B., McCormick, F. and Hawkins, P. T. (1997). Dual role of phosphatidylinositol-3,4,5-trisphosphate in the activation of protein kinase B. Science 277, 567–570.

Sugiyama, M. G., Fairn, G. D. and Antonescu, C. N. (2019). Akt-ing Up Just About Everywhere: Compartment-Specific Akt Activation and Function in Receptor Tyrosine Kinase Signaling. Front Cell Dev Biol 7, 70.

Sun, Y., Thapa, N., Hedman, A. C. and Anderson, R. a (2013). Phosphatidylinositol 4,5-bisphosphate: targeted production and signaling. BioEssaysJ: news and reviews in molecular, cellular and developmental biology 35, 513–22.

Takeuchi, K. and Ito, F. (2011). Receptor tyrosine kinases and targeted cancer therapeutics. Biol Pharm Bull 34, 1774–1780.

Traynor-Kaplan, A., Kruse, M., Dickson, E. J., Dai, G., Vivas, O., Yu, H., Whittington, D. and Hille, B. (2017). Fatty-acyl chain profiles of cellular phosphoinositides. Biochimica et Biophysica Acta - Molecular and Cell Biology of Lipids 1862, 513–522.

Várnai, P. and Balla, T. (1998). Visualization of phosphoinositides that bind pleckstrin homology domains: Calcium- and agonist-induced dynamic changes and relationship to myo-[3H]inositol-labeled phosphoinositide pools. Journal of Cell Biology 143, 501–510.

Wang, S. E., Shin, I., Wu, F. Y., Friedman, D. B. and Arteaga, C. L. (2006). HER2/Neu (ErbB2) signaling to Rac1-Pak1 is temporally and spatially modulated by transforming growth factor beta. Cancer Res 66, 9591–9600.

Wang, C., Faloon, P. W., Tan, Z., Lv, Y., Zhang, P., Ge, Y., Deng, H. and Xiong, J. W. (2007). Mouse lysocardiolipin acyltransferase controls the development of hematopoietic and endothelial lineages during in vitro embryonic stem-cell differentiation. Blood 110, 3601– 3609.

Wills, R. C., Doyle, C. P., Zewe, J. P., Pacheco, J., Hansen, S. D. and Hammond, G. R. V. (2023). A novel homeostatic mechanism tunes PI(4,5)P2-dependent signaling at the plasma membrane. J Cell Sci 136, jcs261494.

Xiong, J. W., Yu, Q., Zhang, J. and Mably, J. D. (2008). An acyltransferase controls the generation of hematopoietic and endothelial lineages in zebrafish. Circulation Research 102, 1057–1064.

Yarden, Y. and Schlessinger, J. (1987). Epidermal growth factor induces rapid, reversible aggregation of the purified epidermal growth factor receptor. Biochemistry 26, 1443– 1451.

Zak, K. and Antonescu, C. N. (2023). Doxycycline-inducible Expression of Proteins at Near-endogenous Levels in Mammalian Cells Using the Sleeping Beauty Transposon System. Bio Protoc 13, e4846.

Zaman, M. F., Nenadic, A., Radojičić, A., Rosado, A. and Beh, C. T. (2020). Sticking With It: ER-PM Membrane Contact Sites as a Coordinating Nexus for Regulating Lipids and Proteins at the Cell Cortex. Frontiers in Cell and Developmental Biology 8,.

Zewe, J. P., Miller, A. M., Sangappa, S., Wills, R. C., Goulden, B. D. and Hammond, G. R. V. (2020). Probing the subcellular distribution of phosphatidylinositol reveals a surprising lack at the plasma membrane. Journal of Cell Biology 219,.

Zhang, K., Chan, V., Botelho, R. J. and Antonescu, C. N. (2023). A tail of their own: regulation of cardiolipin and phosphatidylinositol fatty acyl profile by the acyltransferase LCLAT1. Biochem Soc Trans BST20220603.

Zhu, G., Fan, Z., Ding, M., Zhang, H., Mu, L., Ding, Y., Zhang, Y., Jia, B., Chen, L., Chang, Z., et al. (2015). An EGFR/PI3K/AKT axis promotes accumulation of the Rac1-GEF Tiam1 that is critical in EGFR-driven tumorigenesis. Oncogene 34, 5971–5982.

