## Supplementary figures and images for "The LCLAT1/LYCAT acyltransferase is required for EGF-mediated phosphatidylinositol-3,4,5-trisphosphate generation and Akt signalling"

### Supplemental Figure S1

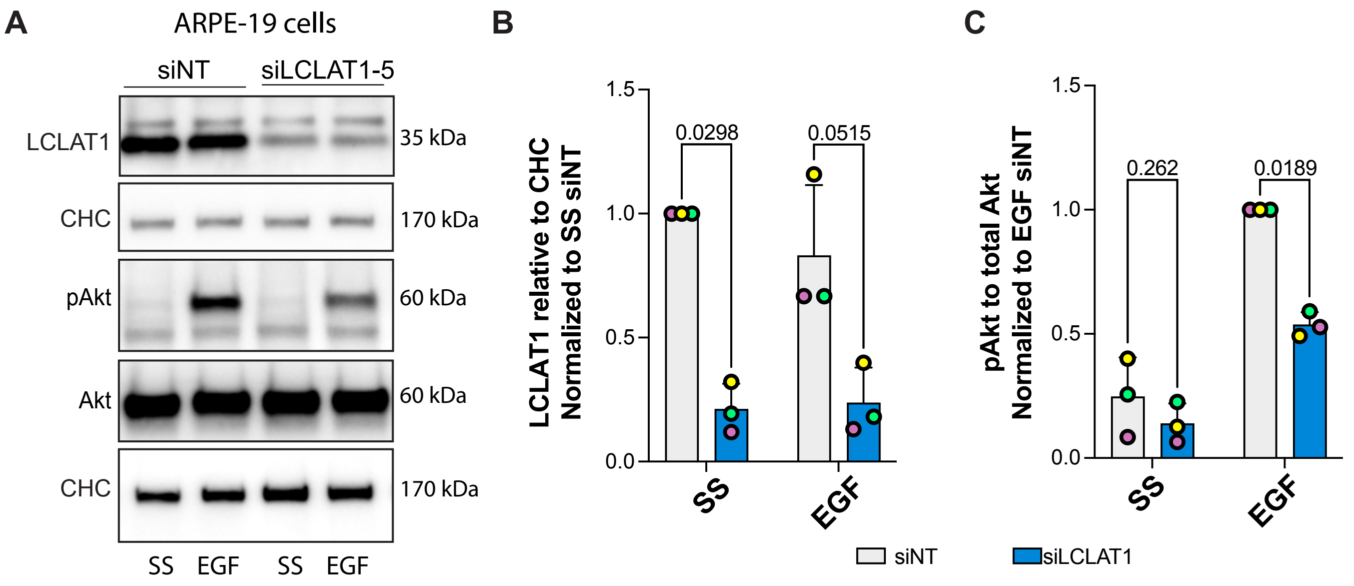


Supplemental Figure S1

### Supplemental Figure S2

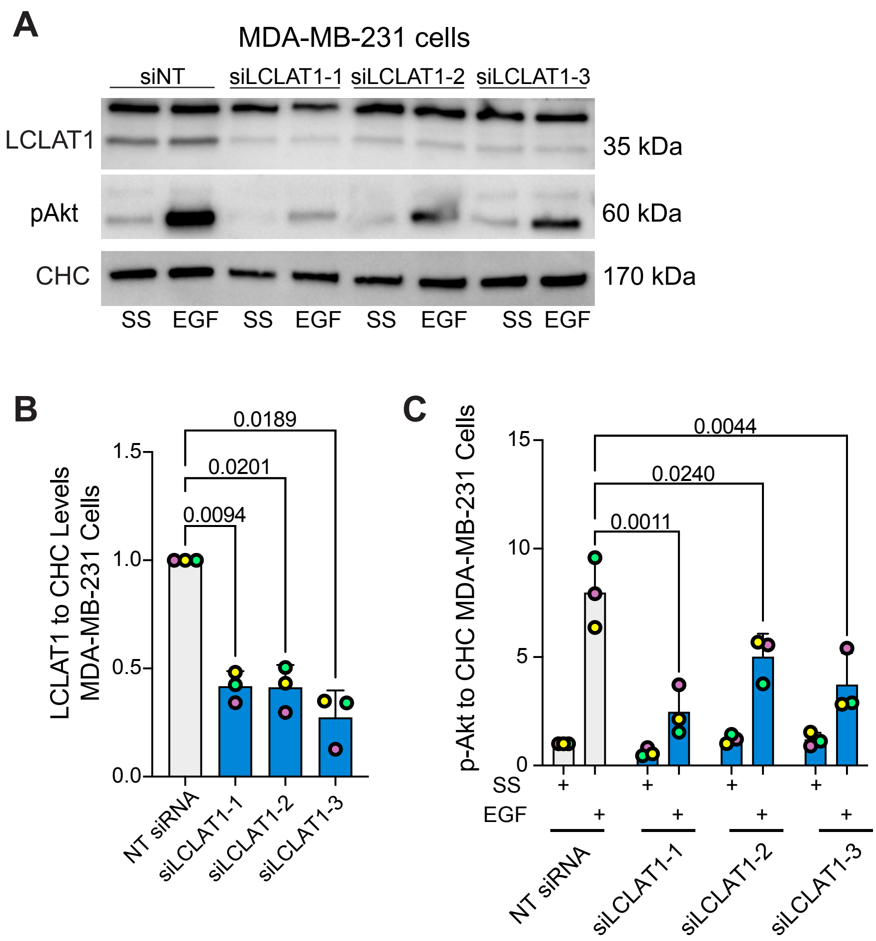


Supplemental Figure S2

### Supplemental Figure S3

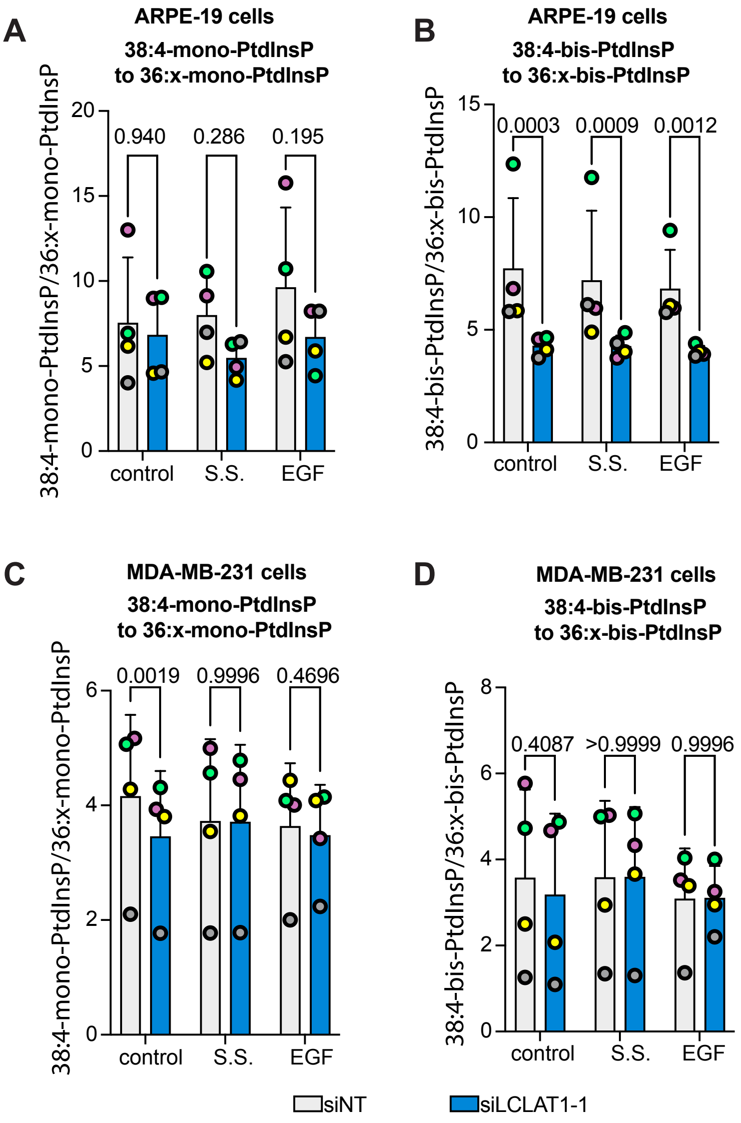


Supplemental Figure S3

### Supplemental Figure S4

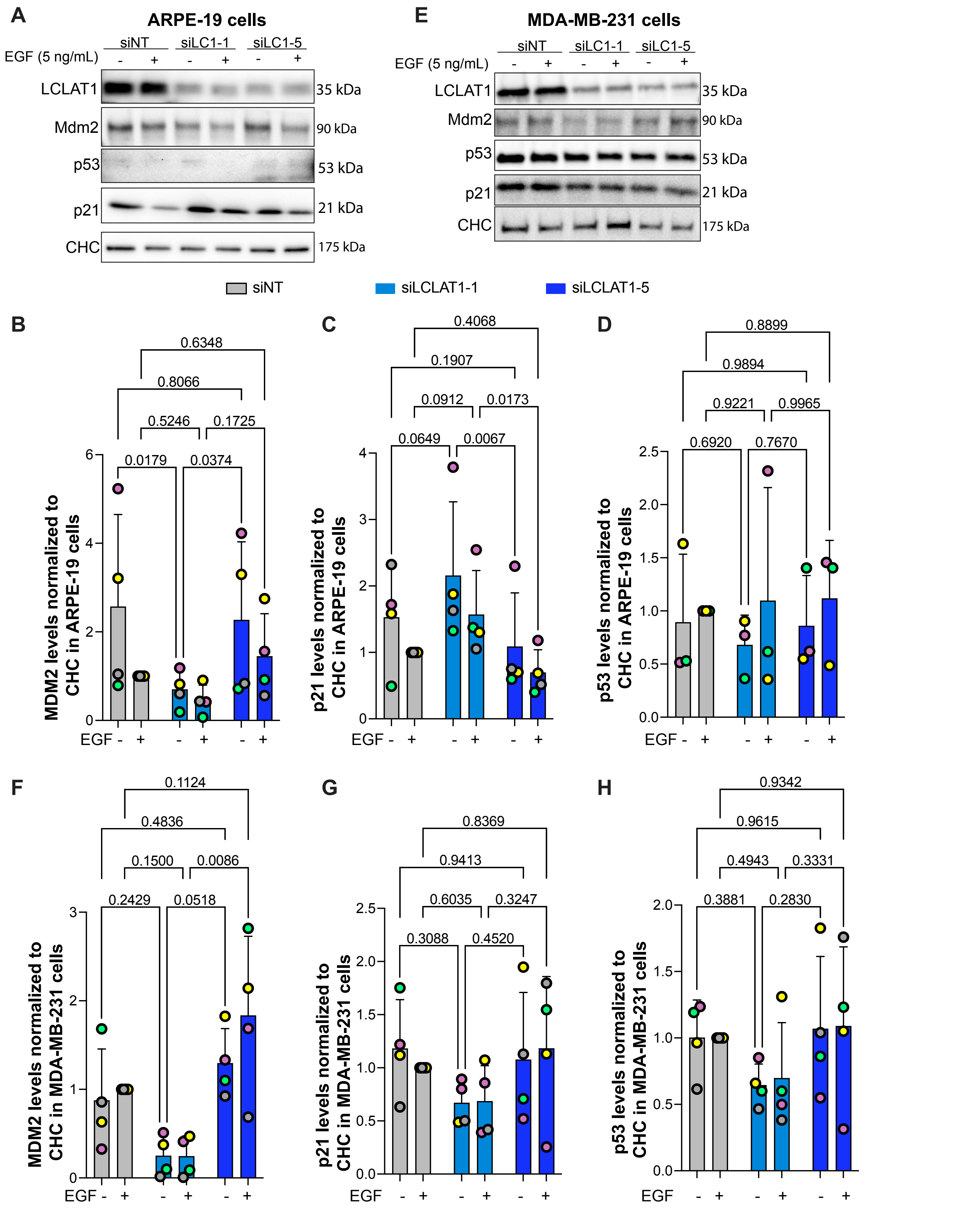


Supplemental Figure S4
